# Disruption of *ADAMTSL4* Causes Ectopia Pupillae in Zebrafish via COL8A1-Driven Cell Migration

**DOI:** 10.1101/2025.11.03.686242

**Authors:** Xinyi Huang, Wannan Jia, Xin Shen, Xinyao Chen, Yalei Wang, Qiuyi Huo, Tianhui Chen, Min Zhang, Keji Jiang, Xuqing Gao, A Guangqi, Fengjing Yang, Yan Pi, Zexu Chen, Yongxiang Jiang

**Affiliations:** State Key Laboratory of Genetic Engineering, National Center for Experimental Biology Education, School of Life Sciences, Fudan University, Shanghai 200438, China; Eye Institute and Department of Ophthalmology, Eye & ENT Hospital, Fudan University, Shanghai 200031, China; Key Laboratory of Myopia and Related Eye Diseases, NHC; Key Laboratory of Myopia and Related Eye Diseases, Chinese Academy of Medical Sciences, Shanghai, 200031, China; East China Sea Fisheries Research Institute, Chinese Academy of Fishery Sciences, Shanghai, Shanghai 200090, China

**Author notes:** Xinyi Huang and Wannan Jia contributed equally to the work presented here and should therefore be regarded as equivalent authors. **Corresponding author:** Yan Pi, MD, PhD, Zexu Chen, MD, PhD, and Yongxiang Jiang MD, PhD, Correspondence: Yan Pi: State Key Laboratory of Genetic Engineering, School of Life Sciences, Fudan University, 2005 Songhu Road, Shanghai 200438, China.; Zexu Chen: Institute and Department of Ophthalmology, Eye & ENT Hospital, Fudan University, 83 Fenyang Road, Shanghai 200031., Yongxiang Jiang: Eye Institute and Department of Ophthalmology, Eye & ENT Hospital, Fudan University, 83 Fenyang Road, Shanghai 200031.

## Abstract

Congenital ectopia lentis (EL) poses a significant threat to visual outcomes in children. Although emerging causative genes have been identified, their biological functions remain poorly understood. In this study, we investigated the contribution of *ADAMTSL4* variants to congenital EL in a cohort of 702 pedigrees. Pathogenic variants in *ADAMTSL4* were identified in 23 probands (3.28%), comprising 29 distinct variants, with 14 being novel, making *ADAMTSL4* the second most frequently mutated gene in the cohort. All affected individuals exhibited EL, and 26.09% also presented with ectopia pupillae (EP), also referred to as ectopia lentis et pupillae. To explore the functional consequences of *ADAMTSL4* deficiency, we generated a CRISPR/Cas9-mediated *adamtsl4*-knockout zebrafish model, that faithfully recapitulated cardinal human disease features with an incidence comparable to that observed in affected patients. Histological and ultrastructural analysis revealed disrupted zonular fiber anchorage at the lens capsule, even in mutated eyes without overt EL or EP. Transgenic overexpression of *adamtsl4* successfully reversed the ocular phenotypes, confirming the gene’s essential role in ocular development. Single-cell RNA-sequencing and fluorescence in situ hybridization demonstrated enriched *ADAMTSL4/adamtsl4* expression in the equatorial lens epithelium, retinal pigment epithelium (RPE), iris anterior pigmented epithelium, and choroid fibroblast. Functional assays using zebrafish and human RPEs revealed that *ADAMTSL4* deficiency compromised cell adhesion and promoted cell migration. Transcriptomic profiling revealed significant enrichment of extracellular matrix organization and cell adhesion pathways, with cross-species validation identifying consistent upregulation of *COL8A1/col8a1b*. Notably, *COL8A1* knockdown in *ADAMTSL4*-deficient RPEs partially reversed the aberrant migratory phenotype, suggesting a functional interaction. Together, these findings establish *ADAMTSL4* as a major causative gene in ectopia lentis et pupillae, highlight its role in orchestrating ocular integrity via regulation of extracellular matrix and cell behavior.

## INTRODUCTION

Congenital ectopia lentis (EL) is the second most prevalent indication for pediatric lens surgery, affecting approximately 6 in 100,000 individuals^[1]^. This condition arises from congenital anomalies causing laxity or disruption of zonular fibers (Zinn’s zonules)^[2]^. Displacement of the crystalline lens from its normal position may lead to significant refractive errors, visual impairment, and form deprivation^[3]^. Although surgical intervention remains the mainstay treatment, procedures carry substantial risks, including suprachoroidal hemorrhage, retinal detachment, and intraocular lens dislocation or subluxation—potentially resulting in permanent vision loss^[4]^. These risks pose considerable burdens on affected families and healthcare systems. Consequently, elucidating the genetic and molecular mechanisms underpinning congenital EL is essential for developing novel therapeutic strategies.

ADAMTSL4, a member of the ADAMTS-like protein family (A Disintegrin And Metalloproteinase with Thrombospondin Motifs-like), is defined by its type I thrombospondin repeats^[5]^. This catalytically inactive secreted glycoprotein localizes to the extracellular matrix (ECM), where it is thought to contribute to ECM remodeling and the maintenance of connective tissue homeostasis^[6]^. Loss-of-function mutations in ADAMTSL4 are associated with a distinct form of congenital EL, frequently co-occurring with ectopia pupillae (EP)—a condition characterized by teardrop-shaped pupils or marked pupillary displacement from the visual axis^[6]^. Given this unique phenotypic combination, the Online Mendelian Inheritance in Man (OMIM) database designates *ADAMTSL4*-associated EL with EP as a distinct disease entity (Ectopia Lentis et Pupillae, OMIM: 225200). EP-induced iris anatomical abnormalities substantially increase surgical complexity in EL cases, predisposing patients to complications such as failed continuous curvilinear capsulorhexis, intraoperative iris hemorrhage, and posterior capsule rupture^[8]^. In severe cases, EP may displace the visual axis, impairing visual signal transmission and disrupting visual development in children, thereby increasing amblyopia risk. Notably, patients with *ADAMTSL4* mutations exhibit significantly poorer visual outcomes and a higher incidence of amblyopia compared to those with *FBN1* mutations, the most common genetic cause of congenital EL..^[9, 10]^ However, the molecular mechanisms underlying EP pathogenesis in *ADAMTSL4* deficiency remain poorly understood, and current *ADAMTSL4* knockout animal models fail to recapitulate the human EP phenotype^[11]^.

Zebrafish have emerged as a powerful model for inherited ocular diseases due to their high genetic homology with humans—approximately 70% of human disease-related genes, including those governing eye development and function, are conserved^[12]^. As diurnal vertebrates, zebrafish share similar ocular architecture and visual processing pathways with humans. Their rapid *ex utero* development and embryonic transparency enable real-time visualization of eye morphogenesis and pathology^[13]^. Moreover, large clutch sizes facilitate high-throughput genetic screening and robust statistical analysis of phenotypic outcomes. In this study, we first identified patients with biallelic *ADAMTSL4* mutations within a large congenital EL cohort and generated *adamtsl4* knockout zebrafish via CRISPR/Cas9 to investigate the molecular mechanisms underlying EP in *ADAMTSL4*-associated congenital EL.

## RESULTS

### Biallelic *ADAMTSL4* Mutations Caused Ectopia Lentis and Ectopia Pupillae in Humans

A total of 702 pedigrees with congenital EL were recruited at the Eye & ENT Hospital of Fudan University between January 2016 and the present. All probands underwent panel-based next-generation sequencing (NGS). Biallelic *ADAMTSL4* mutations were identified in 23 probands (3.28%) (**Figure 1A and Supplementary Table 1**), making *ADAMTSL4* the second most frequently mutated gene in the cohort **(Figure 1B**). Phenotypic analysis revealed that all affected individuals exhibited EL. Among these, about 20% presented with unilateral EL, approximately 20% with asymmetric bilateral EL, and 60% with symmetric bilateral EL. EP was observed in 26.09% of patients, with approximately 30% of those cases involving only one eye. Additional ocular anomalies included microspherophakia (21.74%), persistent pupillary membranes (72.19%), poor pupillary dilation (52.17%), and congenital cataracts (21.74%) (**Figure 1C**). Representative clinical images are shown in **Figure 1D and 1E**. Co-segregation analysis confirmed that the identified *ADAMTSL4* variants followed an autosomal recessive inheritance pattern, with all variants occurring in trans (**Figure 1F**).

**Figure 1.**
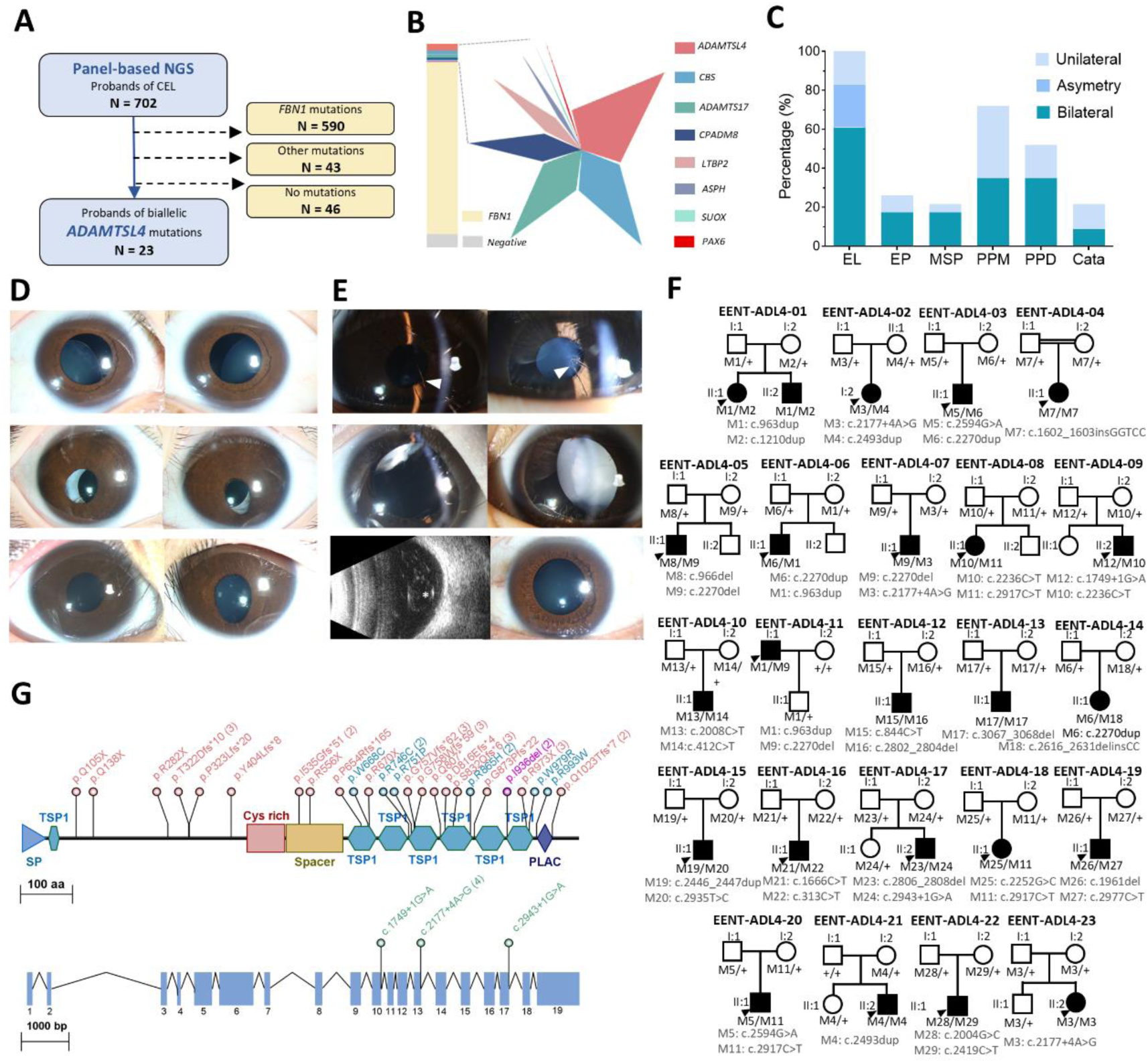
Clinical and Genetic Characterization of Biallelic *ADAMTSL4* Mutations Identified in the Patient Cohort. (A). Schematic representation of the patient recruitment process. A total of 23 probands with biallelic *ADAMTSL4* mutations were identified from a cohort of 702 congenital EL probands. (B). Proportional contribution of each gene associated with congenital EL identified in this cohort. (C). Distribution of ocular phenotypes observed in patients with biallelic *ADAMTSL4* mutations, categorized by unilateral, asymmetrical, and bilateral involvement. (D). Representative clinical features of probands with bilateral inferior-temporal EL (arrows) and PPD (top panel), bilateral inferior-temporal EP (arrowheads) after IOL implantation (middle panel), and unilateral droplet-like EP (arrowhead) with bilateral MSP (bottom panel). (E). Representative clinical features of probands with bilateral PPM (arrowheads) with asymmetrical EL in the right eye (upper panel), bilateral dense CC with asymmetrical EL in the right eye (arrows) (middle panel), and lens dislocation (asterisk) into the vitreous body in the right eye as revealed by B-ultrasound but mild EL in the left eye (arrows) (bottom panel). (F). Pedigree diagrams and segregation analysis of 23 pedigrees with congenital EL and biallelic *ADAMTSL4* mutations. Probands are indicated by arrowheads, and familial segregation patterns are shown to demonstrate inheritance. (G). Overview of the mutational spectrum of *ADAMTSL4* at the genomic and protein levels in the patient cohort. Mutations are color-coded as follows: frameshift (red), missense (cyan), inframe deletion (purple), and splicing (green). Numbers in parentheses indicate the count of probands carrying each specific variant. Scale bar, 100aa (protein); 1000 bp (genome). CC, congenital cataract; EL, ectopia lentis; EP, ectopia pupillae; IOL, intraocular lens; OD, oculus dexter; OS, oculus sinister; PPD, poor pupillary dilation; PPM, persistent pupillary membrane; MSP, microspherophakia.

A total of 29 distinct *ADAMTSL4* variants were identified, 14 of which are novel (**Figure 1G**). All variants were classified as pathogenic or likely pathogenic according to ACMG guidelines (**Supplementary Table 2**). The most common mutation type was frameshift (37.93%), followed by nonsense (24.14%), missense (20.69%), splicing (10.34%), and in-frame deletion (6.90%). All missense variants were subjected to further *in silico* analysis (**Supplementary Figure 1**). Protein sequence alignment revealed that the affected residues were evolutionarily conserved across major vertebrates. Structural modeling further indicated that all but one missense variant were predicted to significantly affect protein stability (|ΔΔG| > 0.5 kcal/mol).

### CRISPR/Cas9-Mediated Knockout of *adamtsl4* Recapitulated Ectopia Lentis and Ectopia Pupillae in Zebrafish

Previous studies reported that N-ethyl-N-nitrosourea (ENU) induced *Adamtsl4* point mutations in mice resulted in EL but failed to reproduce EP, limiting their utility for mechanistic investigation^[11]^. To better model the human ocular phenotype, we employed zebrafish as an alternative model. The zebrafish a*damtsl4* protein is evolutionarily conserved, containing all canonical domains found in vertebrates (**Supplementary Figure 2A**). Sequence alignment revealed 38% overall amino acid identity with the human ADAMTSL4 protein, with individual conserved domains showing 36% to 63% identity (**Supplementary Figure 2B and 2C**).

To elucidate the molecular role of *ADAMTSL4*, we generated *adamtsl4* knockout zebrafish using CRISPR/Cas9-mediated gene editing. A single guide RNA (sgRNA) targeting exon 3 of *adamtsl4* was designed and screened, resulting in three independent mutant lines: *adamtsl4* ^△ 5^, *adamtsl4* ^△ 7^, and *adamtsl4* ^△ 22^ (**Supplementary Figure 3**). All mutations introduced frameshifts within exon 3, leading to premature truncation of the protein (**Figure 2A**). Correspondingly, *adamtsl4* mRNA expression was significantly reduced in all lines (**Figure 2B**). The mutants survived into adulthood. Phenotypic assessment at one month of age revealed EP in approximately 10% to 50% of the individuals across different mutant lines (**Figure 2C**). Affected fish exhibited markedly deviated, undersized pupils, and dislocated lenses situated within the vitreous cavity, rendering the lens undetectable in the anterior chamber (**Figure 2D**) — a phenotype absent in wild-type controls. Although these abnormalities were most pronounced in adults, early signs, including pupil malformation and absence of lens in the anterior chamber, were evident by 7 days post-fertilization (dpf) and progressed through 2 months post-fertilization (mpf) (**Figure 2D**). Subsequent functional studies focused primarily on the *adamtsl4*^△7^ line, hereafter referred to as *adamtsl4^−/−^* for simplicity.

**Figure 2.**
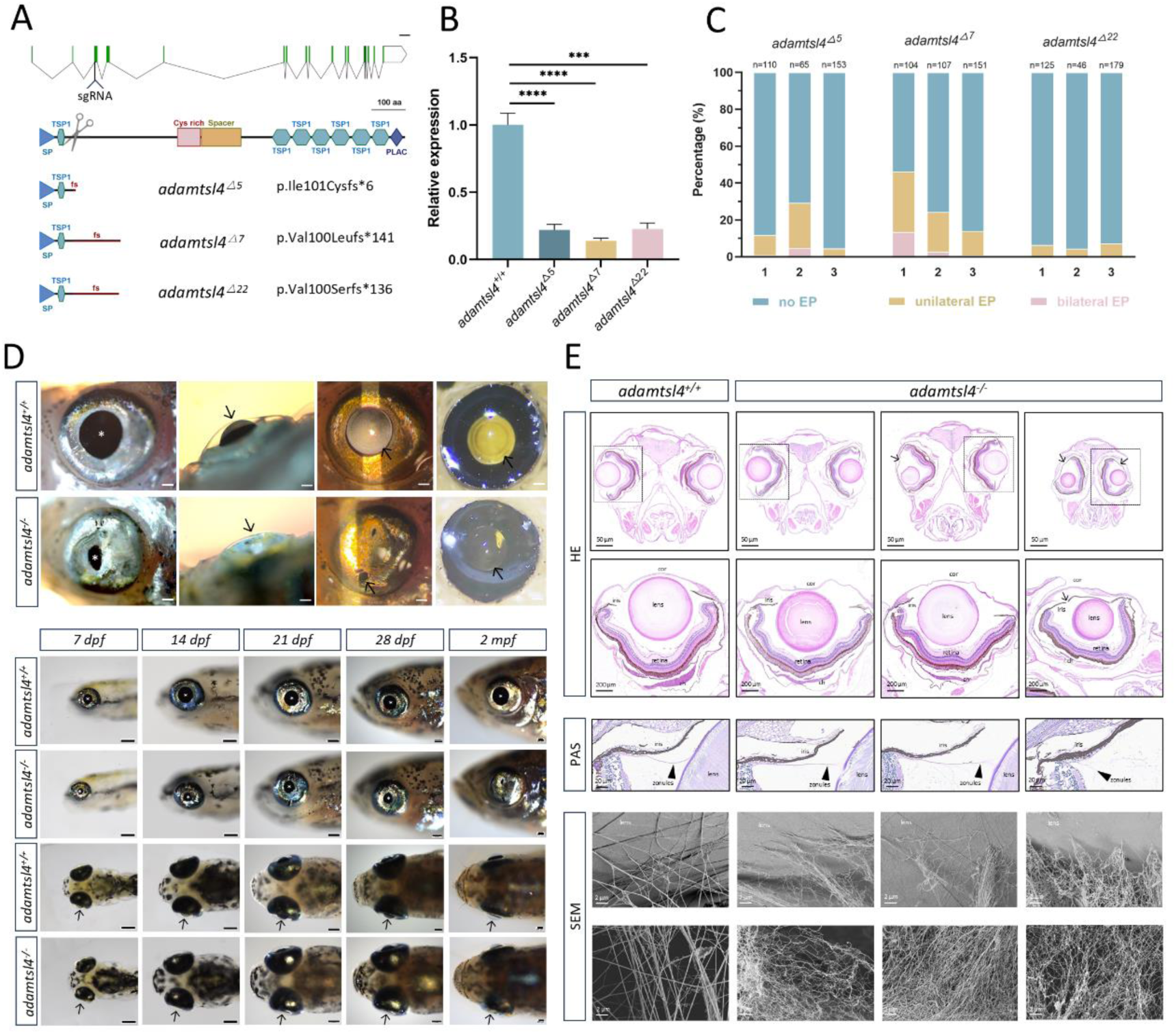
Loss of *adamtsl4* Function Leads to EL and EP in zebrafish. (A). Schematic representation of the CRISPR/Cas9-mediated knockout strategy for *adamtsl4*, illustrating the locations of sgRNA target sites and resulting frameshift variants (*adamtsl4^△5^*, *adamtsl4^△7^*, *adamtsl4^△22^*). Scale bar, 2 kb (genome); 100aa (protein). (B). qPCR analysis showing significant downregulation of *adamtsl4* mRNA expression in all knockout lines compared with sibling controls. (C). Distribution of ocular phenotypes in *adamtsl4^-/-^*mutant lines at 1 mpf: categorized as no EP, unilateral EP, and bilateral EP. Each genotype was analyzed in triplicate. (D). Representative ocular phenotypes of *adamtsl4^-/-^* mutants at 1 mpf. Mutants displayed smaller, decentered pupils (asterisks) and the absence of the lens (arrows) in the anterior chamber. Dissection at the equatorial plane revealed lens (arrows) displacement into the vitreous cavity. Phenotypic progression was traceable as early as 7 dpf, with the smaller pupil (asterisks) and absence of the lens (arrows) becoming increasingly pronounced over time. Scale bars, 200 μm. (E). Histological and ultrastructural comparison of eyes from *adamtsl4^⁻/⁻^* mutants and *adamtsl4^⁺/⁺^* sibling controls at 3 mpf. Panels include: a normal eye in a sibling control (first row), a grossly normal eye from a mutant without EP (second row), a grossly normal eye from a mutant with unilateral EP (third row), and an affected eye from a mutant with bilateral EP (fourth row). EP was indicated by arrows. Boxed areas are shown at higher magnification. PAS staining visualized the zonular fibers (arrowheads), which were broken at the lens end in the EP eye. SEM revealed fragile zonule-lens attachment and sparse, thin zonules even in phenotypically normal-appearing mutant eyes. Statistical significance is indicated by asterisks: ns *P* ≥ 0.05, * *P* < 0.05, ** *P* < 0.01, *** *P* < 0.001, **** *P* < 0.0001; error bars indicate the standard error of the mean. ch, chroid; cor, cornea; dpf: day post-fertilization; EL, ectopia lentis; EP, ectopia pupillae; fs, frameshift; mpf, month post fertilization; PAS, Periodic Acid Schiff; qPCR, quantitative polymerase chain reaction; SEM, Scanning electron microscopy; sgRNA, single guide RNA.

### Zonular Disruption Underlies EP and EL Phenotypes in *adamtsl4^⁻/⁻^* Zebrafish

To further elucidate the pathological changes resulting from *adamtsl4* deficiency, multiple histological analyses were conducted. The Hematoxylin and Eosin (HE) staining of tissue in *adamtls4^-/-^*mutant eyes with EP showed the posterior dislocation of the lens into the vitreous body and significant pupillary displacement (**Figure 2E**, HE panel). Periodic Acid-Schiff (PAS) staining showed disrupted Zinn’s zonules at their attachment site to the lens in affected eyes, suggesting that *adamtsl4* deficiency impairs zonular anchorage at the lens capsule (**Figure 2E**, PAS panel). This finding was further supported by scanning electron microscopy (SEM), which demonstrated zonular breakage near the lens equator, along with disorganized and weakened zonules—even in eyes without overt EL (**Figure 2E**, SEM panel). These observations suggest that, although only ∼30% of *adamtsl4*^⁻/⁻^ mutants exhibited overt EP and EL, zonular integrity was broadly compromised, including in eyes appearing grossly normal. This may reflect an early or subclinical stage of EL. Taken together, these data indicate that *adamtsl4^⁻/⁻^* zebrafish faithfully model the ocular pathologies seen in congenital EL patients with *ADAMTSL4* mutations.

A broader ocular phenotype, extending beyond overt EP and EL, was also assessed. Given that the ‘grossly normal’ eyes of *adamtsl4^-/-^*mutants also exhibited disorganized zonules, these eyes were classified as EL eyes for simplicity (**Supplementary Figure 4A**). Quantitative histological analysis revealed significantly reduced lens thickness in EP eyes of *adamtsl4^-/-^*mutants compared to normal sibling controls (*P* = 0.0192). Central corneal thickness was also significantly decreased in both EL (*P* = 0.0008) and EP (*P* < 0.0001) eyes compared to controls. Additionally, the choroidal area in the median sagittal sections was markedly reduced in EP eyes (*P* = 0.0044) but not in EL eyes (*P* = 0.9350) (**Supplementary Figure 4B**). These findings indicate that *adamtsl4* mutations cause a spectrum of ocular abnormalities in zebrafish, extending beyond EL and EP.

### Overexpression of *adamtsl4* via Tol2 Transposition Rescued the Ocular Anomalies in *adamtsl4* Knockout Mutants

To further validate the specificity of ocular phenotypes associated with *adamtsl4* deficiency, we generated a transgenic zebrafish line, Tg(*hsp:adamtsl4:E2A:EGFP*), in which *adamtsl4* expression was driven by a heat shock promoter (**Supplementary Figure 5A**). Following a 1-hour incubation at 37 °C, *adamtsl4* expression was significantly upregulated. The transgenic line was crossed with the *adamtsl4^-/-^* mutant line, and the resulting progeny were incrossed to generate a stable rescue line. (**Supplementary Figure 5B**). Ocular phenotypes and biometric parameters of 7 dpf larvae were quantified and compared across groups (**Supplementary Figure 5C**). The *adamtsl4^-/-^*mutant line exhibited a significantly increased anterior chamber depth, indicative of posterior lens dislocation, whereas the rescue line demonstrated values comparable to those of sibling controls. Similarly, lens thickness was increased in the *adamtsl4^-/-^*mutants, consistent with a more spherical lens shape resulting from zonular weakness; this phenotype was reversed in the rescue line. Axial length was also significantly elongated in the mutant line but normalized in both the overexpression and rescue groups. The refractive error had no significant differences among groups, likely due to compensatory axial elongation counterbalancing the optical effects of posterior lens displacement and increased lens power. Collectively, these findings demonstrate that adamtsl4 overexpression effectively rescued the early manifestations of EL in the *adamtsl4^-/-^*mutant zebrafish.

### Spatiotemporal Expression Patterns of *ADAMTSL4/adamtsl4* in Zebrafish and Human Ocular Tissues

To further elucidate the molecular function of *ADAMTSL4/adamtsl4*, we examined its expression patterns in both zebrafish and human tissues. In zebrafish larvae, *adamtsl4* mRNA levels showed a progressive increase during development (**Figure 3A**). In adult zebrafish, adamtsl4 was highly enriched in the testis, eye, heart, and skeletal muscle, as revealed by tissue-specific expression profiling (**Figure 3B**). Given this broad extraocular expression, we assessed systemic phenotypes in *adamtsl4^-/-^*mutants (**Supplementary Figure 6**). Notably, body length was significantly reduced in both unilateral (*P* = 0.0030) and bilateral EP mutants (*P* < 0.0001), but not in bilateral EL mutants (*P* = 0.2445), when compared with sibling controls (**Supplementary Figure 6A**). Despite the high expression of *adamtsl4* in cardiac tissues, histological analysis revealed no significant abnormalities in the cardiac muscle or aortic wall of mutant zebrafish (**Supplementary Figure 6B**).

**Figure 3.**
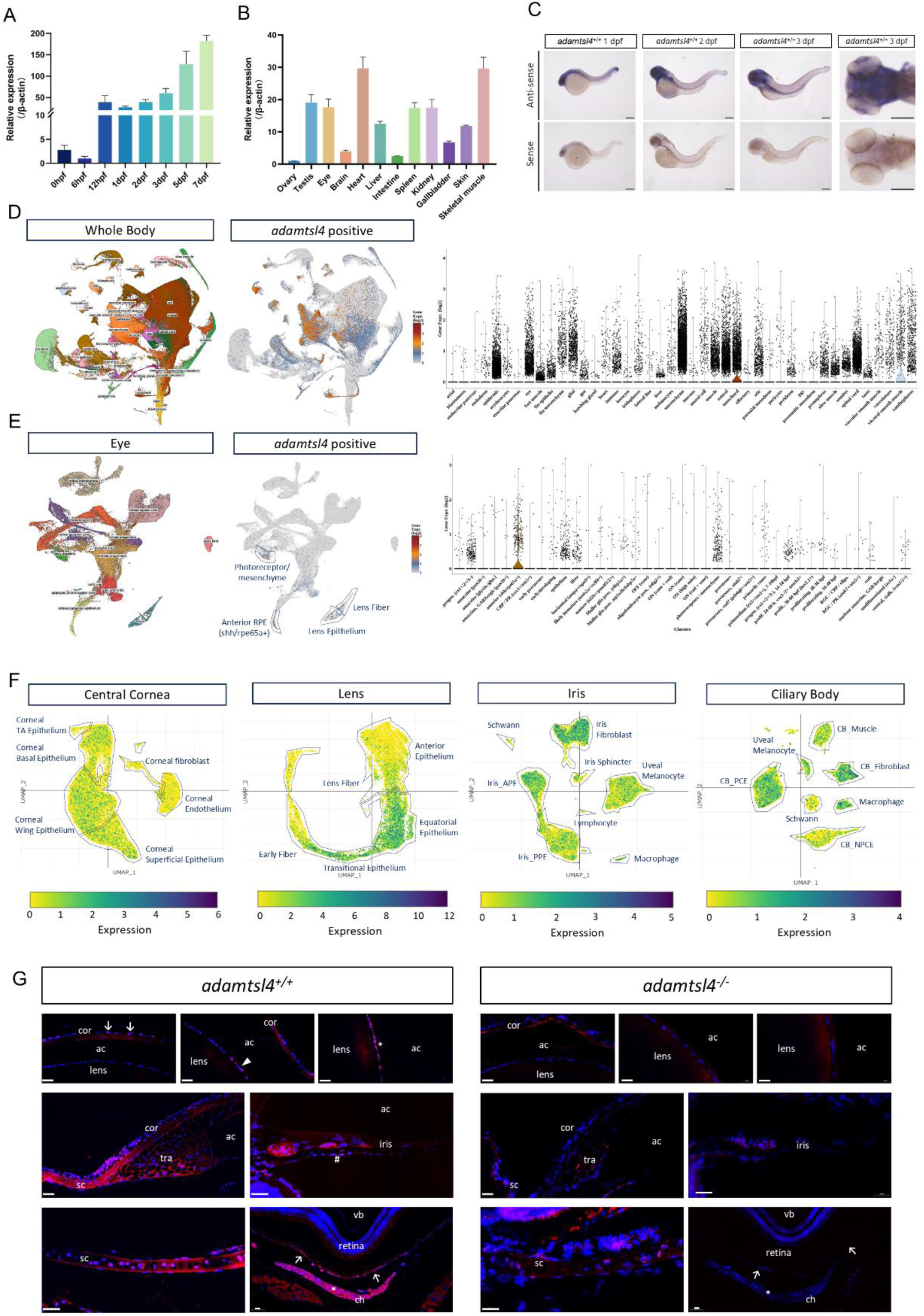
Spatiotemporal Expression Profile of *ADAMTSL4/adamtsl4* in Zebrafish Larvae and Human Ocular Tissues. (A). Developmental expression of *adamtsl4* during early zebrafish larval stages, assessed by qPCR. β-actin was used as an internal control, and expression levels were normalized to the developmental stage with the lowest expression. (B). Tissue-specific expression of *adamtsl4* in adult zebrafish, analyzed by qPCR. β-actin served as an internal reference gene, and the tissue with the lowest expression level was used for normalization. (C). Whole-mount in situ hybridization of *adamtsl4* mRNA in wild-type larva at 1 dpf, 2 dpf, and 3 dpf, showing *adamtsl4* expression in the brain, notochord, and ocular regions. A clear contrast between antisense and sense probe signals confirmed probe specificity. Scale bar: 200 μm. (D). UMAP embedding of scRNA-seq data from whole zebrafish larvae merged across developmental stages (48–120 hpf), illustrating *adamtsl4* expression levels by color intensity and violin plots grouped by tissue types. (E). UMAP embedding of scRNA-seq data from extracted ocular tissue of larva (48–120 hpf), highlighting *adamtsl4* expression patterns by cell type via color mapping and violin plots. (F). UMAP plot of human anterior eye scRNA-seq data, with cells colored by *ADAMTSL4* expression and boxed according to cell clusters. (G). Fluorescent in situ hybridization of adult zebrafish eyes at 3 mpf using an antisense *adamtsl4* probe. *adamtsl4* mRNA was detected in the central corneal epithelium (arrows), anterior lens epithelium near the equatorial (arrowhead), equatorial lens epithelium (asterisk), trabecular meshwork cells, iris stroma cells and epithelium (pound signs), scleral stroma cells, RPE (arrows), and choroidal stroma (asterisks). DAPI counterstaining marked nuclei (blue). Probe specificity was validated using eyes from *adamtsl4^⁻/⁻^* mutant zebrafish. Scale bars: 20 μm. Error bars in the bar plot indicate the standard error of the mean; the vertical line in the violin graph indicates the distribution range. ac, anterior chamber; APE, anterior pigmented epithelium; CB, ciliary body; ch, chroid; cor, cornea; dpf: day post-fertilization; hpf, hour post-fertilization; NPCE, non-pigmented ciliary epithelium; PGC, primordial germ cell; prec., precursor; progen., progenitor; PCE, pigmented ciliary epithelium; PPE, posterior pigmented epithelium; qPCR, quantitative polymerase chain reaction; RBC, retinal bipolar cell; RGC, retinal ganglion cells; RPE, retinal pigmented epithelium; mpf, month post fertilization; sc, sclera; scRNA-seq, single-cell RNA sequencing; tra, trabecular meshwork; UMAP, uniform manifold approximation and projection; vc, vitreous cavity.

The spatial expression pattern of adamtsl4 was examined from 1 to 7 days dpf using whole-mount in situ hybridization, revealing prominent signals in the brain, eye, and notochord (**Figure 3C**). The single-cell RNA-seq (scRNA-seq) atlas of zebrafish larva further showed high expression of *adamtsl4* in xanthophores, vasculature, mesenchyme, epidermis, eye, and notochord (**Figure 3D**). The *adamtsl4* showed increased expression in visceral smooth muscle but maintained a steady expression level in the ocular system (**Supplementary Figure 7A**). The expression of *adamtsl4* in the ocular system larval was highly enriched in certain regions, including the photoreceptor, anterior retinal pigmented epithelium (RPE), and lens (**Figure 3E**). Among the *adamtsl4*-expressing cells, the *shh/rpe65a*-positive RPE was of particular interest, as the *adamtsl4* showed the most abundant expression and the increased tendency during the development (**Supplementary Figure 7B and 7C)**. The scRNA-seq atlas of the human anterior eye showed a more detailed expression of *ADAMTLS4* in the cornea, lens, iris, and ciliary body (**Figure 3F**). *ADAMTSL4* is selectively expressed in the equatorial lens cells, transitional lens epithelium, and subsets of early lens fiber, but showed relatively low expression in the anterior lens epithelium. In the iris, *ADAMTSL4* was expressed across all major cellular clusters, with notable enrichment in the anterior pigmented epithelium (APE) and fibroblasts. The pigmented ciliary epithelium (PCE), which is anatomically continuous with the APE, exhibited a similarly high level of *ADAMTSL4* expression, as did fibroblasts within the ciliary body. In contrast, the posterior pigmented epithelium (PPE) of the iris and the non-pigmented ciliary epithelium (NPCE), which are also structurally aligned, showed relatively lower *ADAMTSL4* expression. The fluorescence *in situ* hybridization was applied to verify the above single-cell expression analysis in zebrafish eye sections and the specificity of the probe was verified by the absence of signal in these tissues in *adamtsl4^-/-^* mutants. (**Figure 3G**). Expression was confirmed in the equatorial lens epithelium, trabecular meshwork, iris, RPE, and choroid. Consistent with the scRNA-seq data, expression was absent from the anterior lens surface but enriched in the lens equatorial cells. The corneal epithelium, sclera, and trabecular meshwork also showed a high expression of *ADAMTSL4*. In the posterior segment of the eye, the *ADAMSTL4* was mainly expressed in the RPE and choroid, which was consistent with the scatter and low expression in the human retina (**Supplementary Figure 7D**) and high expression in the choroid fibroblasts (**Supplementary Figure 7E**)

### *ADAMTSL4* Deficiency Impairs Cell Adhesion and Enhances Migration Without Affecting Viability

To further elucidate the molecular function of *ADAMTSL4*, we examined cellular alterations following its downregulation in human cells. A cultured human RPE cell line (ARPE-19) was selected as the experimental model, based on the following rationale: the robust expression of adamtsl4 in the anterior RPE of zebrafish larvae, the developmental and anatomical continuity among the iris APE, the ciliary body PCE, and the retinal RPE, and the practical availability and suitability of human RPE cells for in vitro manipulation. Efficient knockdown of *ADAMTSL4* mRNA expression was achieved using a validated small interfering RNA (siRNA) (**Figure 4A**). Ki67 immunostaining indicated no significant difference in proliferative capacity between *ADAMTSL4* knockdown cells and control cells (**Figure 4B**). Though the EdU-positive cells slightly increased in the *ADAMTSL4* knockdown cells (**Figure 4C**). Apoptotic rates remained unchanged, as demonstrated by both Annexin V staining (**Figure 4D**) and TUNEL assays (**Figure 4E**). These findings suggested the dispensability of *ADAMTSL* in maintaining cell viability in RPEs.

**Figure 4.**
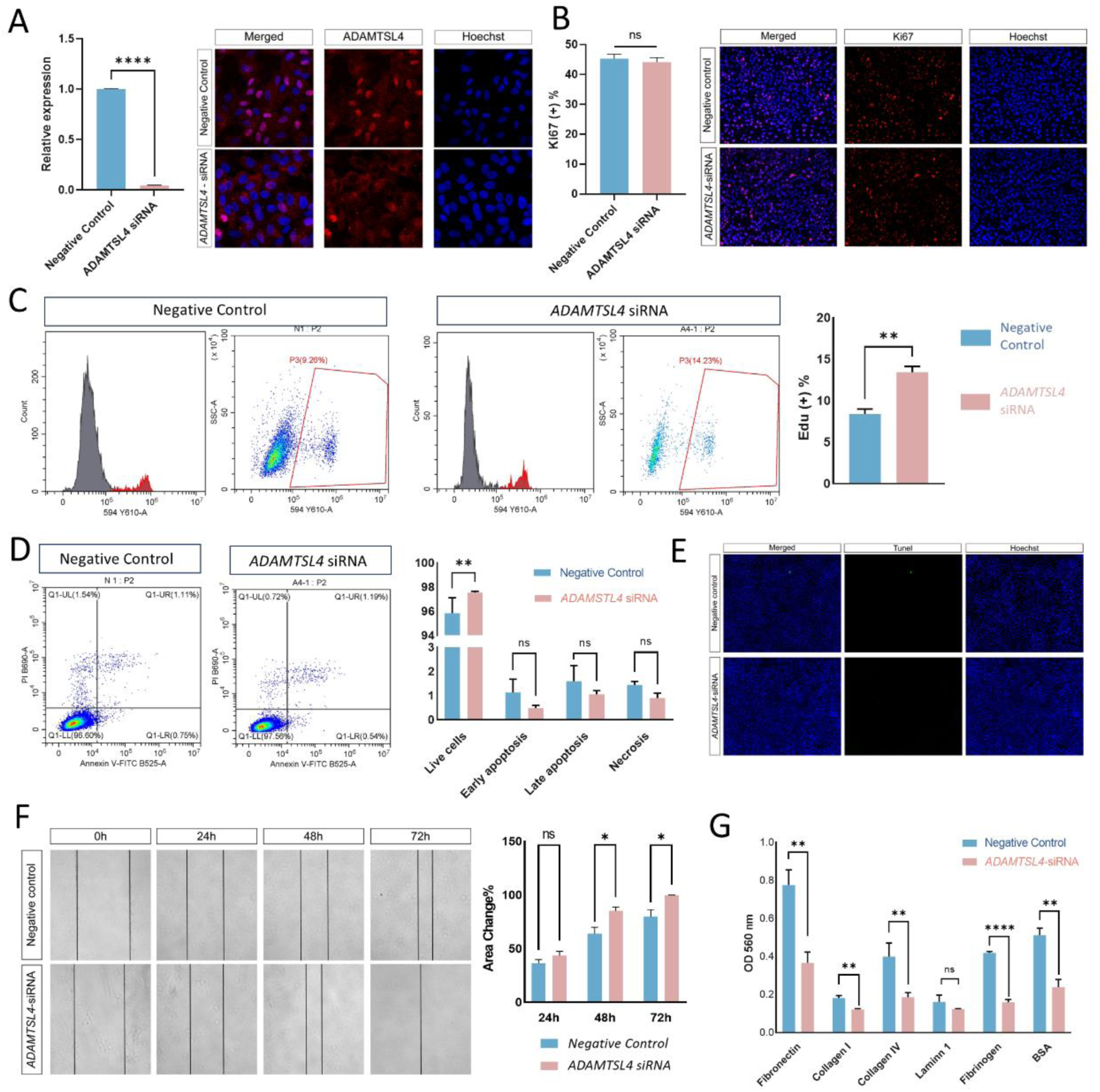
*ADAMTSL4* Deficiency Impairs Cell Adhesion and Promotes Migration Without Altering Proliferation or Apoptosis in Human RPEs. (A). qPCR and immunofluorescence analysis confirmed a significant reduction in *ADAMTSL4* mRNA and protein levels in RPEs following siRNA-mediated knockdown, compared with negative siRNA controls. (B). Proliferation was assessed by quantifying Ki67-positive nuclei, expressed as a percentage of total DAPI-stained cells in control and *ADAMTSL4* knockdown RPEs. (C). Assessment of cell proliferation by EdU incorporation assay followed by flow cytometry analysis. Representative flow cytometry plots and quantitative data are shown for each group. (D). Assessment of cell apoptosis by Annexin V/PI staining and flow cytometry. Cells were classified as live (Annexin V⁻/PI⁻), early apoptotic (Annexin V⁺/PI⁻), late apoptotic (Annexin V⁺/PI⁺), or necrotic (Annexin V⁻/PI⁺). Representative dot plots and quantitative analysis are shown for each group. (E). Representative images of TUNEL-stained indicated no change of cell apoptosis level in *ADAMTSL4* knockdown RPEs. (F). Wound healing assays demonstrated significantly enhanced migratory capacity in *ADAMTSL4* knockdown RPEs. Wound closure was quantified as the percentage of area covered after incubation, across at least three biological replicates per group. (G). Quantification of ECM-mediated cell adhesion using a colorimetric assay. Cells were seeded onto 48-well plates pre-coated with fibronectin, collagen I, collagen IV, laminin I, fibrinogen, or BSA. The adherent cells were fixed, stained, and quantified. Statistical significance is indicated by asterisks: ns *P* ≥ 0.05, * *P* < 0.05, ** *P* < 0.01, *** *P* < 0.001, **** *P* < 0.0001; error bars indicate the standard error of the mean. BSA, Bovine Serum Albumin; ECM, extracellular matrix; OD, optical density; qPCR, quantitative polymerase chain reaction; RPE, retinal pigmented epithelium; siRNA, small interfering RNA.

Strikingly, cell migration was significantly enhanced in *ADAMTSL4* knockdown RPEs, as shown by wound healing assays (**Figure 4F**). Concordantly, cell adhesion was markedly reduced across most ECM coatings in knockdown cells (**Figure 4G**). Together, these results indicate that while *ADAMTSL4* does not appear to regulate cell proliferation or apoptosis, it plays a critical role in modulating cell adhesion and migration.

### RNA-seq Analysis of *adamtsl4*-Regulated Genes and Pathways

To investigate the downstream molecular targets regulated by *adamtsl4* that may contribute to the EP phenotype, we performed bulk RNA-seq analysis on EP eyes from *adamtsl4^-/-^*zebrafish and normal eyes from wild-type sibling controls. The clear separation between groups along the principal components suggests a differential transcriptome profile (**Figure 5A**). Differential expression analysis identified 1,672 upregulated and 552 downregulated genes in the *adamtsl4^-/-^*group (**Figure 5B and 5C**). Gene Ontology (GO) and Kyoto Encyclopedia of Genes and Genomes (KEGG) enrichment analyses revealed significant alterations in categories related to cell adhesion (GO:0007155, GO:0022610, GO:0007156, GO:0098609; KEGG: dre04514, dre04510), extracellular matrix organization (GO:0005576, GO:0044421, GO:0005615), and calcium signaling pathways (GO:0005509; KEGG: dre04020) (**Figure 5D and 5E**). Gene Set Enrichment Analysis (GSEA) further confirmed the enrichment of ECM-related pathways (**Figure 5F**), highlighting a key role for *adamtsl4* in modulating ECM composition and organization.

**Figure 5.**
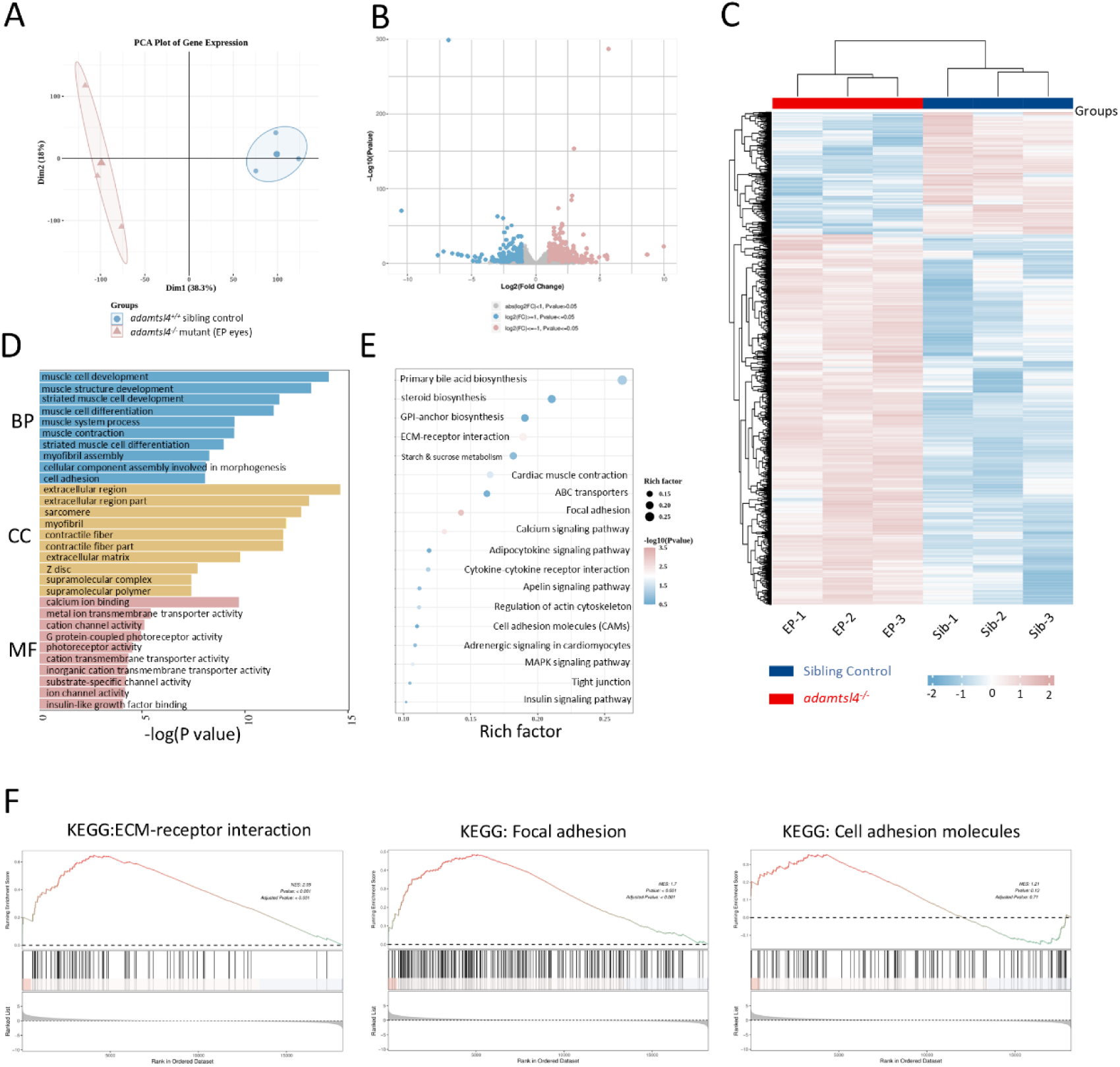
Transcriptomic Alterations in EP Eyes of *adamtsl4^⁻/⁻^* Mutant Zebrafish Revealed by RNA-seq Analysis. (A). PCA plot of RNA-seq samples revealed clear transcriptomic separation between EP eyes of *adamtsl4⁻/⁻* mutants (red) and normal eyes of sibling controls (blue), indicating substantial gene expression divergence. (B). Volcano plot shows differentially expressed genes between the EP eye of *adamtsl4^-/-^*mutant with the normal eye of the sibling control. Each point represents a gene, with the X-axis of log_2_(fold change) in gene expression and the Y-axis representing –log₁₀(*P* value). Red and blue dots denote upregulated and down-regulated genes, respectively (|log_2_(Fold change)| ≥ 1). Gray dots indicate no significant DEGs. (C). Heatmap of DEGs between EP mutant and control eyes. Rows represent genes, and columns represent individual samples from either control (blue) or *adamtsl4⁻/⁻* EP eyes (red). (D). GO enrichment analysis showing the top 10 significantly enriched terms from the BP, MF, and CC categories. The X-axis indicates –log₁₀(*P* value), and the Y-axis lists enriched GO terms. (E). Significantly Enriched KEGG Pathways. Dot plot showing the top 20 significantly enriched KEGG pathways based on candidate gene set enrichment analysis. The X-axis represents the rich factor. The Y-axis depicts –log₁₀(*P* value). The size of the bubbles represents the gene count for each pathway. Pathways are ranked by rich factors in descending order. (F). GSEA analysis for ECM-related highlighting coordinated transcriptional changes in ECM components in *adamtsl4^-/-^* mutant zebrafish eyeballs. Statistical significance is indicated by asterisks: ns *P* ≥ 0.05, * *P* < 0.05, ** *P* < 0.01, *** *P* < 0.001, **** *P* < 0.0001; error bars indicate the standard error of the mean. BP, Biological Process; CC, Cellular Component; DEG, differentially expressed gene; EP, ectopia pupillae; ECM, extracellular matrix; FC, fold change; FDR, false discovery rate-adjusted *P* value; GO, Gene Ontology; GSEA, Gene Set Enrichment Analysis; KEGG, Kyoto Encyclopedia of Genes and Genomes; MF, Molecular Function; NES, normalized enrichment score; PCA, principal component analysis.

### Disruption of *ADAMTSL4/Adamtsl4* Upregulated *COL8A1*/*Col8a1b* Expression

To identify downstream targets regulated by *adamtsl4*, we conducted quantitative PCR (qPCR) analysis on 20 representative ECM-related genes (**Figure 6A**). Consistent with the RNA-seq data, several ECM-related genes, including *lgals1l1*, *col6a1*, *col6a2*, *col8a1b*, *col112a1a*, *tgfb2*, *fbn1*, *fbn2*, *frem2b*, *tem176*, were significantly upregulated in *adamtsl4^-/-^*zebrafish, whereas genes such as *efnb2a* and *lim2.2* were markedly downregulated. To assess cross-species relevance, we evaluated the expression of orthologous genes in human RPEs following *ADAMTSL4* knockdown. Among these, only *COL8A1* and *TGFB2* exhibited consistent upregulation (**Figure 6B**). We then examined their protein expression levels via western blotting and found that only COL8A1 was significantly increased in *ADAMTSL4*-deficient cells (**Figure 6C)**. Though, co-immunoprecipitation assays revealed no direct interaction between ADAMTSL4 and COL8A1 (**Figure 6D**). To investigate the functional consequence of *COL8A1* upregulation, we knocked down *COL8A1* in *ADAMTSL4*-deficient RPEs (**Figure 6E**). This combinatorial knockdown reversed the enhanced migratory phenotype observed in *ADAMTSL4*-deficient RPEs (**Figure 6F**). Collectively, these findings suggested that the loss of *ADAMTSL4/adamtsl4* led to upregulation of *COL8A1/col8a1b*, which in turn contributed to increased cell migration. The main findings of this study are summarized in **Figure 7**.

**Figure 6.**
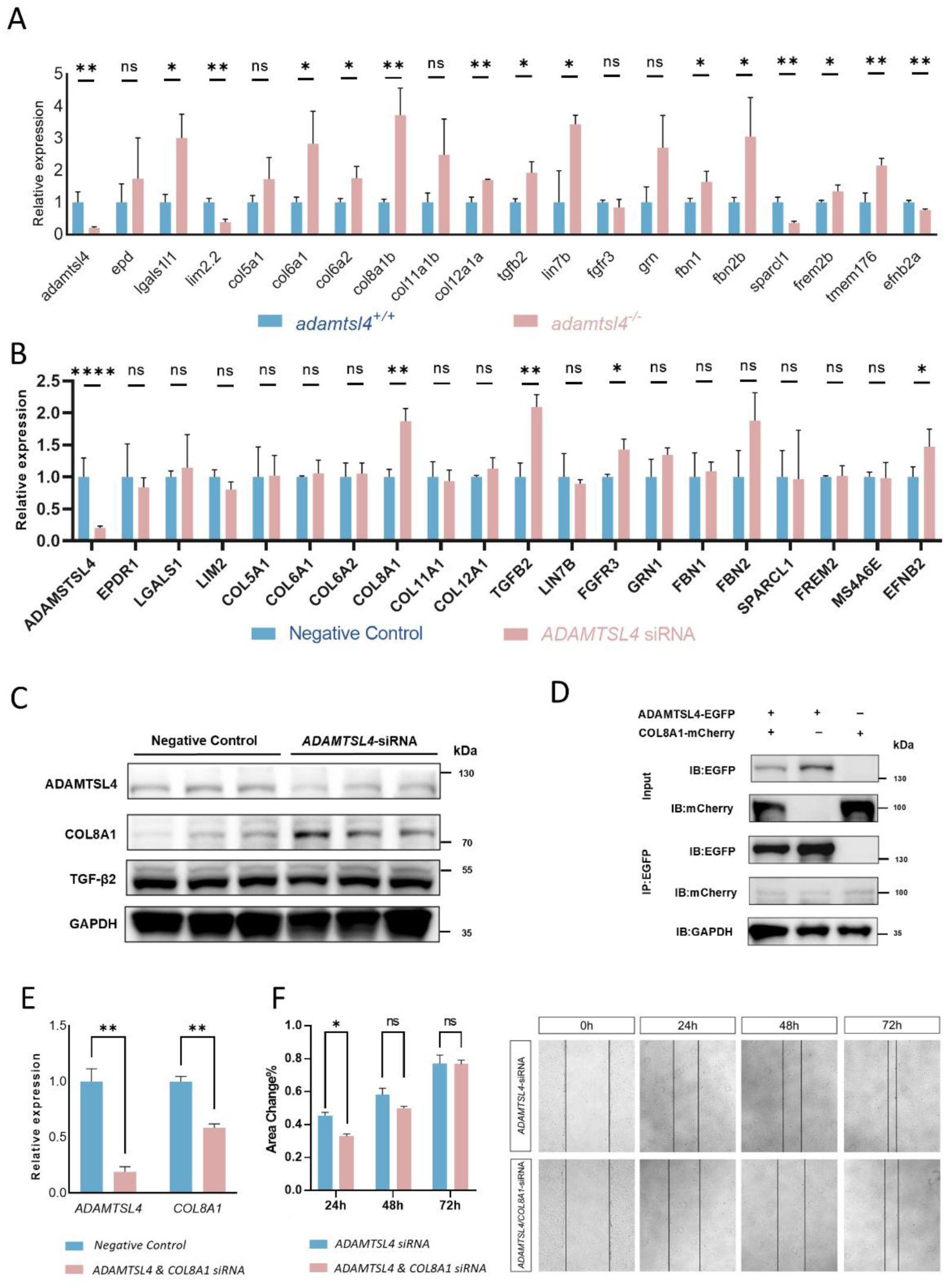
Loss of *ADAMTSL4/adamtsl4* Leads to Upregulation of *COL8A1* which Alters Cell Migration. (A). qPCR analysis showing fold changes in mRNA expression of ECM-associated genes in EP eyes of *adamtsl4^⁻/⁻^* zebrafish mutants compared with normal eyes of sibling controls. Each group included three biological replicates. (B). qPCR validation of ECM-associated gene expression in human RPEs following siRNA-mediated knockdown of *ADAMTSL4*, compared to negative siRNA controls. Each group has three biological replications. (C). Western blot analysis of ADAMTSL4, TGFB2, and COL8A1 protein levels in *ADAMTSL4* knockdown RPEs versus controls. GAPDH was used as a loading control. Each group contained three biological replicates. (D). Co-IP between ADAMTSL4 and COL8A1 in HEK293T cells. Exogenous *ADAMTSL4* fused with *EGFP* and *COL8A1* fused with mCherry were expressed and precipitated using EGFP antibody-coated beads. Interacting proteins were detected by Western blot. (E). Knockdown of *COL8A1* in *ADAMTSL4-*deficient RPE cells using siRNA. Knockdown efficiency was confirmed by qPCR. (F). Wound healing assay showing significantly reduced cell migration in RPEs following *ADAMTSL4* knockdown, *COL8A1* knockdown, combined knockdown. Negative siRNA control was transfected toaccount for potential cytotoxic effects of transfection, and total siRNA concentrations were maintained consistently across all groups. Quantification was based on the percentage change in wound area across at least three independent experiments. Statistical significance is indicated by asterisks: ns *P* ≥ 0.05, * *P* < 0.05, ** *P* < 0.01, *** *P* < 0.001, **** *P* < 0.0001; error bars indicate the standard error of the mean. Co-IP, Co-Immunoprecipitation; qPCR, quantitative polymerase chain reaction; siRNA, small interfering RNA; RPE, retinal pigmented epithelium; HEK293T, Human Embryonic Kidney 293T cell; IP, immunoprecipitation; IB, immunoblotting.

**Figure 7.**
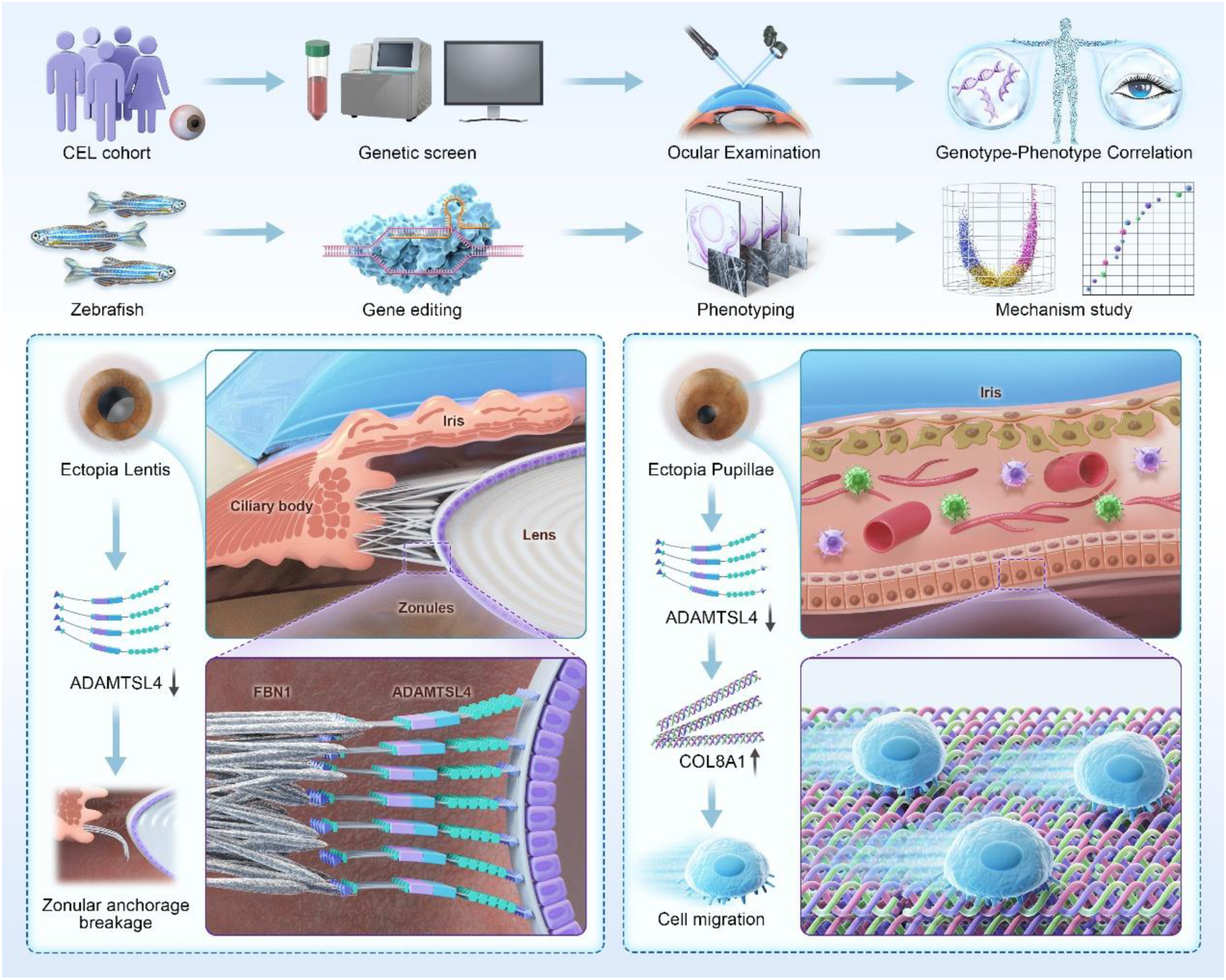
Overview of the Study Design and Key Conclusions. This study was prompted by clinical genetic screening and genotype-phenotype correlation analysis in patient cohort with congenital EL. We employed an integrative, multidisciplinary approach -combining clinical genomics, vertebrate modeling, single-cell transcriptomics, and functional cellular assays -to investigate the role of ADAMTSL4 in EL and EP. Our findings establish ADAMTSL4 as a principal causative gene for EL with co-occurring EP, define its spatial and temporal expression patterns, and elucidate its dual role in maintaining zonular fiber anchorage at the lens equator and restraining aberrant cell migration during iris and pupil formation. EL, ectopia lentis; EP, ectopia pupillae.

## DISCUSSION

Genetic mutations represent the primary pathogenic driver of congenital EL, with causative variants identifiable in over 80% of affected individuals^[14]^. Advances in next-generation sequencing have substantially expanded the genetic landscape of congenital EL^[15, 16]^. Although most of these genes encode components of the ECM or its regulators, their precise biological roles remain poorly understood. Mechanistic insight has been limited, in part due to the lack of reliable animal models that faithfully replicate this highly specialized ocular phenotype. This study provides compelling genetic and functional evidence that biallelic mutations in *ADAMTSL4* represent one of the most prevalent causes of congenital EL, frequently accompanied by EP. By integrating comprehensive genotypic screening in a large clinical cohort with functional studies in a CRISPR/Cas9-engineered zebrafish model, we uncovered essential roles for *ADAMTSL4* in maintaining zonular fiber integrity and regulating cell migration during ocular development. These findings not only established a robust vertebrate model that faithfully recapitulated the human phenotype but also expanded our understanding of the molecular basis of *ADAMTSL4*-associated ocular disease.

The co-occurrence of EP strongly suggests the presence of biallelic *ADAMTSL4* mutations in patients with congenital EL. However, previous studies using ENU-induced *Adamtsl4* missense mutant mice successfully recapitulated EL but failed to model EP, thereby limiting their utility in mechanistic studies^[11]^. In contrast, zebrafish offer unique advantages for modeling anterior segment disorders, including high optical transparency, proportionally large eyes, conserved ocular anatomy, and genetic tractability^[13, 17]^. We demonstrated that *adamtsl4* knockout in zebrafish leads to early-onset, progressive, and reproducible core phenotypes of this disease identity—including EL and EP—with an incidence comparable to that observed in affected patients. The phenotypic robustness, despite lower amino acid identity between zebrafish and human ADAMTSL4 compared to murine orthologs, likely reflects the complete loss of function induced by frameshift alleles. This is consistent with clinical genotype-phenotype correlations, in which patients carrying truncating variants exhibit a higher prevalence of EP compared to those with missense ones^[18, 19]^. Moreover, *adamtsl4* overexpression fully rescued ocular abnormalities in mutants, underscoring the gene’s causal role in observed ocular anomalies.

Mechanistically, ADAMTSL4 lacks the N-terminal metalloprotease domain found in canonical ADAMTS family members but retains multiple glycosaminoglycan-binding and thrombospondin type 1 repeat (TSR) domains^[6]^, suggesting its function in ECM organization or molecular interaction. While the function of *ADMTSL4* remained largely unknown, three mechanistic hypotheses emerged from current evidence. First, ADAMTSL4 may modulate ADAMTS protease activity via shared C-terminal auxiliary domains, similar to its homolog papilin, which inhibits ADAMTS2 activity and cooperates with MIG-17/ADAMTS in nematode development^[20, 21]^. Second, it may mediate ECM–cell interactions, anchoring zonular fibers to the lens capsule ^[22]^, analogous to TSR-mediated pericardial cell attachment in C. elegans^[23]^. Supporting this model, we observed disrupted zonular attachment to the lens capsule in *adamtsl4* knockout zebrafish, and reduced cell adhesion with increased migration in *ADAMTSL4*-deficient human RPE cells. Third, ADAMTSL4 likely regulates ECM synthesis and deposition. Both ADAMTSL4 and its structural homolog ADAMTSL6 enhance FBN1 deposition^[24]^, with *ADAMTSL6* overexpression partially rescuing phenotypes in *Fbn1* mutant mice^[25]^. Though the *ADAMTSL4* deficiency did not induce significant fold change of *FBN1* or *FBN2* expression in RPE, suggesting alternative downstream targets.

We propose a tissue-specific model of *ADAMTSL4* function in the ocular system: in the lens equator, ADAMTSL4 maintains zonular anchorage; while in the iris and ciliary body, it modulates migratory behavior of iris progenitor cells that would alter the pupillary positioning. Although the precise mechanism underlying EP remains unresolved, our data support a model in which enhanced cellular motility contributes to ectopic pupil positioning. The iris - a circular pigmented structure positioned between the cornea and lens -comprises multiple layered components: fibroblasts, melanocytes, vascularized stroma, APE, and PPE layers^[26]^. In zebrafish embryos, we observed that *adamtsl4* expression is enriched in anterior RPE marked by *shh* and *rpe65a*. Notably, in the human eye, *ADAMTSL4* is also highly expressed in the iris APE and ciliary body PCE—structures that are anatomically continuous with, and embryologically derived from, the anterior RPE^[27, 28]^. During embryogenesis, the epithelial components of the iris originate from the peripheral rim of the optic cup, whereas the stromal tissue derives from the periocular mesenchyme^[29]^. Critical to this process, neural crest cells migrate along epithelial surfaces to differentiate into iris stroma, corneal stroma, and trabecular meshwork, with precise cellular localization being essential for normal iris formation^[30]^. Disruption of cellular positioning at this stage—through enhanced migration or failed adhesion—may lead to asymmetric pupil formation or EP. This theory parallels congenital aniridia, in which *PAX6* mutations impair epithelial differentiation ^[31, 32]^; and Axenfeld-Rieger syndrome, where *PITX2* and *FOXC1* mutations compromise periocular mesenchymal migration^[33, 34]^, resulting in a spectrum of iris anomalies including EP, polycoria, and stromal hypoplasia^[35]^.

Further reinforcing this model, we identified *COL8A1* as a conserved downstream effector of *ADAMTSL4*. Upregulated in both *adamtsl4^−/−^* zebrafish and *ADAMTSL4*-deficient human RPEs, *COL8A1* promoted cell motility through integrin-mediated focal adhesion or ECM remodeling^[36–38]^, which aligned with previous literature showing *COL8A1*-driven migration enhancement across multiple cancer types^[39–41]^. Knockdown of *COL8A1* reversed the migratory phenotype in *ADAMTSL4*-deficient RPEs, establishing a functional link. Although no direct interaction was observed between ADAMTSL4 and COL8A1, these findings suggest that ADAMTSL4 regulates COL8A1 expression indirectly to restrain excessive cell migration and to ensure proper iris and pupil formation. The discovery of this axis provides a novel mechanistic link between ECM remodeling and iris morphogenesis.

## LIMITATION

Several limitations should be acknowledged when interpreting these findings. First, although our zebrafish model faithfully recapitulates both EL and EP, interspecies differences —such as the higher incidence of unilateral EP in zebrafish—may reflect species-specific modifiers or developmental plasticity. Second, the mechanism by which ADAMTSL4 regulates COL8A1 expression remains incompletely defined, as no direct physical interaction was detected. Future studies will be required to elucidate the transcriptional or post-transcriptional pathways linking these molecules. Third, incomplete penetrance of EP in both patients and animal models suggests that additional genetic or epigenetic modifiers contribute to the phenotype. Future identification of these modifiers may uncover mechanisms of disease resilience and inform therapeutic targeting.

## CONCLUSION

This study identifies biallelic mutations in *ADAMTSL4* as the second major genetic contributor to congenital EL and identifies *ADAMTSL4* as a pleiotropic regulator of ocular ECM integrity and anterior segment morphogenesis. Through the establishment of a genetically tractable zebrafish model and the elucidation of a novel ADAMTSL4–COL8A1 regulatory axis, we provide critical mechanistic insight into the development of EL and EP. These findings lay the groundwork for further exploration of ECM-targeted therapies in *ADAMTSL4*-associated ocular disorders.

## METHODS

### Patient Cohort and Genetic Sequencing

The study was approved by the Human Research Ethics Committee of the Eye & ENT Hospital of Fudan University (ChiCTR2000039132) and conducted in accordance with the principles of the Declaration of Helsinki. Written informed consent was obtained from all participants or legal guardians for those under 18 years old. The recruitment strategy for the congenital EL cohort has been described in detail in our previous publication^[14]^. All participants underwent comprehensive ophthalmic evaluations, and detailed medical and family histories were recorded. Panel-based NGS designed by Amplicon Gene (Shanghai, China), as previously reported^[42, 43]^. In brief, Genomic DNA was extracted from peripheral blood samples and sequenced using the Illumina NovaSeq 6000 platform (Illumina Inc., San Diego, CA, USA). Putative pathogenic variants were validated by Sanger sequencing. When available, co-segregation analysis was conducted on family members using the same approach.

### *In silico* Analysis of the Candidate Variants

The reference sequence of the *ADAMTSL4* transcript was NM_019032. Allele frequency annotation (gnomAD), splicing site prediction (SpliceAI), and pathogenicity prediction (SIFT, PolyPhen, and MutationTaster) were conducted using integrated *in silico* algorithms, the Ensembl Variant Effect Predictor 105 (http://uswest.ensembl.org/info/docs/tools/vep/index.html). All variants were graded according to the American College of Medical Genetics and Genomics guidelines^[44]^. All the mutations were mapped to protein and genomic structures with the IBS1.0.3 illustrator^[45]^. The protein domains were annotated by MOTIF Search (https://www.genome.jp/tools/motif/). Phylogenetic tree, evolutionary analyses, and protein sequence alignment were conducted via MEGA7 software^[46]^. The full-length wild-type protein structure was obtained using AlphaFold, and regions with low prediction confidence were removed using PyMOL. The resulting structure was used as the initial model for all-atom molecular dynamics simulations, which were performed using the AMBER 24 software package. The alteration of the protein stability was indicated by the free energy change value (ΔΔG, kcal/mol). Values greater than 0.5 kcal/mol imply a large increase in protein stability, and values less than −0.5 kcal/mol indicate a large decrease in protein stability.

### Zebrafish Maintenance and Strains

The zebrafish were raised and maintained in accordance with a standard protocol. The wild-type zebrafish are exclusively AB strain in this study. The *adamtsl4* mutant was generated using the CRISPR/Cas9 technology. The designed sgRNA (TCTGACTGCAGAGATCACCG) was transcribed from the linearized pMD19-T-gRNA vector (CZRC, China) *in vitro*. The *Cas9* protein and *adamtsl4* sgRNA were co-injected into the one-cell stage embryos. The knockout efficiency was tested using Sanger sequencing by the detection of overlapping chromatogram peaks. The positive founders were mated with the wild-type fish to obtain F1 generation. The F1 heterozygous zebrafish with identical frameshift mutations were intercrossed to generate an F2 homozygous mutant. F2 homozygous mutants were intercrossed for several generations to obtain maternal-effect-free homozygous mutants for subsequent phenotypic analysis. The primers used for genotyping were: TTATTCTTGACAGGCAGCAGGAC and GACCAATCCCACTCACCTTCTG.

The Tg (*hsp:adamtsl4:E2A:EGFP*) were generated via the Tol2 transposition system. The cDNA fragment of adamtsl4 was cloned into the pT2KhspGFF (CZRC, China). The transposase mRNA was generated by *in vitro* transcription with a linearized plasmid pCS-zTP. A Tol2-donor plasmid DNA and the transposase mRNA were introduced into the zebrafish fertilized eggs by microinjection. The positive founders were mated with wild-type fish to obtain F1 generation. The larvae were raised at 29 ℃ and heat-shocked for 1 h before screening or experiments. In the rescue experiment, *adamtsl4^-/-^*mutant zebrafish were crossed with the Tg (hsp:adamtsl4:E2A:EGFP) line. Fluorescent-positive offspring were identified through heat-shock induction to generate the rescue F1 generation. The F1 fish were then intercrossed, and fluorescent-positive F2 progeny were screened by heat shock. The adult fish of F2 were genotyped to identify *adamtsl4^-/-^*individuals. These F2 mutants were subsequently intercrossed to obtain the rescue F3 generation. Relevant phenotypic data were then collected to assess whether the transgene expression successfully rescued the mutant phenotype.

### Ocular Measurement and Morphological Analysis

In vivo imaging of zebrafish larvae and adults was performed using a stereoscopic fluorescence microscope (M205FA, Leica, Germany) to capture lateral and vertical views. Before imaging, the zebrafish were anesthetized with tricaine (A5040, Sigma). Larvae were immobilized with methyl cellulose (HY-125861, MCE) and treated with phenylthiourea (P7629, Sigma) to inhibit pigmentation when necessary. Body length, anterior chamber depth, axial length, lens thickness, and corneal curvature radius were measured using ImageJ (64-bit Java 1.8.0_172). The refraction error of the larva eye was calculated according to the previous study^[47]^.

### Transcriptome Profiling and Analysis

Eyeballs were collected from 3 mpf wild-type siblings and *adamtsl4^-/-^* mutant zebrafish. For each sample, four eyeballs were pooled per tube, and three biological replicates were prepared per group. After rinsing with PBS, the eyeballs were transferred to pre-chilled 1.5 ml RNase-free microcentrifuge tubes, which were wrapped in aluminum foil and snap-frozen in liquid nitrogen. Samples were then immediately stored at −80 °C. High-throughput sequencing of the above samples was performed using paired-end sequencing on the Illumina HiSeq platform (Shanghai Xuran Biotechnology Co., Ltd.). Adapter sequences and low-quality fragments were dynamically trimmed from the 3’ ends of the sequencing data using Skewer, and quality control analysis of the preprocessed data was performed using FastQC. Differentially expressed genes between sample groups were identified using the DESeq2 package. Genes with |log₂FC| ≥ 1 and *P*-value ≤ 0.05 were considered significantly differentially expressed. GO and KEGG enrichment analyses were performed on the differentially expressed genes using the clusterProfiler R package. GO terms and KEGG pathways with adjusted P-values (Benjamini–Hochberg correction) less than 0.05 were considered significantly enriched. Rich Factor was calculated as the ratio of the number of genes from the candidate gene set annotated to a given pathway to the total number of genes annotated to that pathway in the background set. GSEA was conducted through the Bioconductor package ‘clusterProfiler’(22455463). The Normalized Enrichment Score (NES) is calculated by dividing the enrichment score (ES) by the mean of the ES values from per mutation of the same gene set to account for gene set size and variability.

### ScRNA-seq Analysis

ScRNA-seq data from the published datasets of the human anterior eye, human posterior eye, and developing zebrafish larvae were analyzed to identify cell-type expression of *ADAMTSL4/adamstl4* using standard bioinformatics pipelines^[48–50]^. scRNA-seq expression values were log-transformed and centered using the mean expression values. Principal Component Analysis (PCA) followed by t-distributed Stochastic Neighbor Embedding (t-SNE) or Uniform Manifold Approximation and Projection (UMAP) to visualize the cellular clusters. The average centered expression values of adamtsl4 were calculated for each cell. Cells were grouped into cell clusters and the relative expression level of a given cell cluster was calculated by a scale function. All data processing and analysis were performed using R (version 4.0) and the ‘Seurat’ package.

### Cell Culture and siRNA Knock Down

The human RPE cell line (ARPE-19, QuiCell, #A125) (RRID:CVCL_IQ60) was obtained from Shanghai QuiCell Biotechnology Co., Ltd. The identity of the ARPE-19 cell line was verified via short tandem repeat (STR) analysis, and no evidence of contamination or misidentification was detected. The cell line was cultured using DMEM/F12 (Gibco, #11320033) + 10% FBS (Gibco, #10099141C) at 37 °C in a humidified incubator with 5% CO₂. Cell morphology was monitored daily, and the culture medium was refreshed every three days. When cell confluency reached over 95%, cells were either passaged or cryopreserved. The siRNA (target sequence: GGACCGTCTTTCGATATAA) was synthesized by Guangzhou RiboBio Co., Ltd. The siRNA was transfected into the RPE under 30%-50% confluency in 6-well plates using Lipofectamine™ RNAiMAX (Invitrogen, #13778150). The knockdown efficiency was tested 48h post the transfection by qPCR using the primers: (ACCTGGAGAAACCCTCACCT and GTCTGTCTTTACAGTTCTCATCTTC).

### Cell Apoptosis and Proliferation Assays

TUNEL staining was applied for evaluating cell apoptosis: RPEs were seeded on glass coverslips in 12-well plates and cultured to ∼80% confluence. Cells were fixed with 4% paraformaldehyde for 30 min, permeabilized with 0.3% Triton X-100 for 5 min, and washed with PBS. TUNEL staining was performed using a commercial kit (Beyotime, #C1086) by incubating cells with 100 μL of detection solution (TdT enzyme and fluorescent label) at 37 °C in the dark for 60 min. After washing, nuclei were counterstained with 1 μg/mL DAPI (Beyotime, #C1002) for 5 min in the dark. Cells were visualized under a fluorescence microscope (excitation 450–500 nm).

Apoptosis was also assessed using Annexin V-FITC/propidium iodide (PI) double staining followed by flow cytometry. Cells were harvested by removing the culture medium and washing adherent cells once with PBS. Cells were then detached using trypsin-EDTA, centrifuged at 1000 × g for 5 minutes, and resuspended in PBS for cell counting. Approximately 5 × 10^4^ to 1 × 10^5^ cells were collected by centrifugation and resuspended in 195 µL of Annexin V binding buffer (Beyotime, #C1062S). 5 µL of Annexin V-FITC and 10 µL of PI were added to each sample. The cells were gently mixed and incubated for 10–20 minutes at room temperature in the dark. During incubation, the cells were resuspended 2–3 times to enhance staining uniformity. After incubation, the samples were placed on ice and immediately analyzed using a flow cytometer.

EdU Incorporation was performed to evaluate cell proliferation: RPE cells at 50–60% confluence in 6-well plates were incubated with 50 μM EdU for 2 hours. Cells were then harvested by trypsinization, centrifuged, and fixed in 4% paraformaldehyde for 15 min at room temperature. After permeabilization with 0.3% Triton X-100, EdU staining was performed using the Click-iT reaction cocktail provided in the Elabscience EdU Assay Kit (Elabscience, #E-CK-A377) according to the manufacturer’s instructions. After the final washes, cells were immediately analyzed by flow cytometry. Proliferating cells were quantified as EdU-positive populations.

### Wound Healing and Cell Adhesion Assay

A wound-healing assay was performed to assess cell migration. Cells were seeded into 6-well plates at a density of (3–5) × 10⁵ cells per well in 2 ml of complete medium containing 10% FBS and incubated for 48–72 hours until a confluent monolayer was formed. A straight scratch was then made across the cell layer using a 200 μl pipette tip, ensuring uniform scratch width. The position of each scratch was marked on the outside of the well to facilitate consistent imaging. Detached cells were removed by washing twice with 1 ml PBS, followed by the addition of 2 ml serum-free medium to each well. The plates were returned to the incubator, and images of the scratch area were captured at 0, 24, 48, and 72 hours using an inverted microscope. Scratch width and area were measured over time to quantify the extent of cell migration.

Cell adhesion was evaluated using CytoSelect™ 48-Well Cell Adhesion Assay (CELL BIOLABS, INC., #CBA-070). Under sterile conditions, the was equilibrated to room temperature for 10 minutes. The cell suspension was prepared at a concentration of 0.1–1.0 × 10⁶ cells/ml in serum-free medium and then was added (150μL) to the wells of ECM Adhesion Plate (48 wells). The Plate was incubated in a cell culture incubator for 30–90 minutes to allow cell adhesion. Following incubation, the medium was aspirated and gently washed 4–5 times with 250 μL of PBS. After removing PBS, 200 μL of Cell Stain Solution was added to each well and incubated at room temperature for 10 minutes. The stain was then washed 4–5 times with 500 μL of deionized water. After the final wash, 200 μL of Extraction Solution was added to each air-dried well, which was incubated on an orbital shaker for 10 minutes. The extracted solution from each well was transferred to a 96-well microplate, and the absorbance was measured at 560 nm using a microplate reader.

### Fluorescence *in situ* Hybridization

Fluorescence in situ hybridization was performed on paraffin sections of adult wild-type siblings and *adamtsl4*⁻/⁻ mutant zebrafish at 3mpf. Tissues were fixed using *in situ* hybridization fixative for at least 12 hours, followed by paraffin embedding and sectioning.

Slides were placed in citrate buffer (pH 6.0) and incubated in a 90 °C water bath for 48 minutes, then treated with 20 μg/ml proteinase K at 40 °C for 10 minutes. Sections were rinsed once with distilled water and three times with PBS (5 minutes each). Pre-hybridization was performed with a pre-hybridization buffer at 37 °C for 1 hour. After removing the buffer, hybridization solution containing probes was applied, and sections were incubated overnight at 40 °C. Sections were washed sequentially with 2× SSC at 37 °C for 10 minutes, 1× SSC at 37 °C for 5 minutes (twice), and 0.5× SSC at room temperature for 10 minutes. For branched DNA probe hybridization, 60 μL of preheated hybridization buffer was added to each slide and incubated in a humidified chamber at 40 °C for 45 minutes, with 50 ml of 2× SSC placed at the bottom of the chamber to prevent drying. Slides were then sequentially washed with pre-warmed 2× SSC, 1× SSC, 0.5× SSC, and 0.1× SSC at 40 °C for 5 minutes each. The sense and anti-sense probes were transcribed from the cloned zebrafish *adamtsl4* cDNA.Signal probe hybridization was performed using probe solution diluted 1:200, incubated at 42 °C for 3 hours, followed by washes with 2× SSC (10 minutes at 37 °C), 1× SSC (twice for 5 minutes), and 0.5× SSC (10 minutes). Nuclei were counterstained with DAPI in the dark for 8 minutes, washed with distilled water, and mounted using an anti-fade mounting medium. Finally, sections were examined and imaged under a fluorescence microscope.

### Immunohistochemistry

For cell immunofluorescence, cells were seeded onto glass coverslips in a 12-well plate and cultured until they reached 80% confluence. The plate was then removed from the incubator, and each well was washed three times with 1 ml pre-warmed (37°C) PBS for 5 minutes per wash. Cells were fixed with 4% paraformaldehyde (PFA) at room temperature for 20 minutes, followed by three washes with PBS. To increase membrane permeability, 0.3% Triton X-100 solution was added to each well and incubated at room temperature for 10 minutes, after which the solution was removed, and the cells were washed again with PBS. Blocking was performed by adding 10% goat serum to each well for 40 minutes at room temperature. After blocking, the blocking solution was discarded, and the primary antibody diluted in 1% bovine serum albumin was added to each well. The plate was incubated overnight at 4°C in a humidified chamber. The following day, the plate was incubated at room temperature for 30 minutes in the dark, and cells were washed three times with PBS for 10 minutes each. The secondary antibody was added to each well and incubated for 1 hour in the dark. The plate was then washed three times with PBS, and 1 μg/ml DAPI staining solution was added to each well, incubating for 5 minutes in the dark. After washing with PBS, the coverslips were carefully removed from the wells, and excess liquid at the edges was blotted with filter paper. A drop of the anti-fade mounting medium was applied to a glass slide, and the coverslip was inverted and placed onto the slide, ensuring even coverage of the anti-fade reagent. Finally, the edges of the coverslip were sealed with nail polish, and the samples were stored at 4°C in the dark. The samples were subsequently observed and imaged using an inverted fluorescence microscope or a confocal microscope. The primary antibodies used in this study were as follows: ADAMTSL4 Rabbit pAb (Abclonal, #A4785, 1:1000 dilution) and Ki67 Rabbit pAb (Abclonal, #A11390, 1:1000 dilution), The secondary antibodies used in this study were as follows: Goat anti-Rabbit IgG (H+L) Alexa Fluor™ Plus 488 (Invitrogen, #A32731TR, 1:1000 dilution) and Goat anti-Mouse IgG (H+L) Alexa Fluor™ Plus 555 (Invitrogen, #A32727, 1:1000 dilution).

### Scanning Electron Microscopy

The eyeballs were dissected from 3-month-old wild-type and *adamtsl4^⁻/⁻^* zebrafish and rinsed three times with PBS. Samples were fixed in 2.5% glutaraldehyde at room temperature in the dark for 2 hours, followed by overnight fixation at 4 °C. The eyeballs were then bisected at the equator, rinsed three times with PBS, and further fixed in 1% osmium tetroxide at room temperature in the dark for 2 hours. Dehydration was performed through a graded ethanol series (30%, 50%, 70%, 80%, 90%, 95%, and twice in 100% ethanol) with each step lasting 15 minutes, followed by a final step in isoamyl acetate for 15 minutes. Samples were then subjected to critical point drying. For conductivity, specimens were mounted onto conductive carbon tape and coated with gold using an ion sputter coater. Finally, the prepared samples were imaged and documented using an ultra-high-resolution scanning electron microscope.

### Immunoblotting and Immunoprecipitation

Immunoprecipitation was performed in HEK293T cells (QuiCell, #2937-500) (RRID:CVCL_0063), and was authenticated by STR profiling and confirmed to be free of cross-contamination. HEK293T cells were chosen for Co-IP due to their unparalleled transient transfection efficiency, SV40 T-antigen-driven protein overexpression, and mammalian post-translational modifications machinery. HEK293T cells were transfected using the pCMV6 plasmids, into which cDNAs of *ADAMTSL4* and *COL8A1* plus indicated tags were cloned, and Lipofectamine 3000 (Invitrogen, # L3000001) was used. Adherent cells were washed twice with ice-cold PBS, scraped, and lysed in lysis buffer supplemented with protease and phosphatase inhibitors (20–30 μL per 1×10⁵ cells) on ice for 30 min with intermittent mixing. Lysates were cleared by centrifugation at 12,000 × g for 10 min at 4 °C, and supernatants were collected. For immunoprecipitation, 500 μL of lysate was incubated with 25 μL of pre-equilibrated GFP-Trap agarose beads (chromotek, #gta) at 4 °C for 1 h with gentle rotation. Beads were washed three times with cold lysis buffer and transferred to new tubes. Bound proteins were eluted by adding 1× SDS sample buffer, followed by heating at 100 °C for 10 min to dissociate immune complexes from the beads. Supernatants were collected after centrifugation at 2,500 × g for 2 min and subjected to SDS–PAGE and immunoblotting.

For immunoblotting, total protein lysates were prepared by lysing cells in 4 mL of RIPA buffer supplemented with 50× protease and phosphatase inhibitors (80 μL each), followed by ultrasonic disruption. After centrifugation at 12,000 × g for 15 min at 4 °C, supernatants were collected, and protein concentrations were determined using the BCA assay. Equal amounts of protein (5–15 μg) were mixed with 5× SDS-PAGE loading buffer, boiled at 100 °C for 10 min, and separated on SDS-PAGE precast gels. Proteins were transferred to PVDF membranes at 280 mA for 90 min. Membranes were blocked for 15 min at room temperature, incubated with primary antibodies overnight at 4 °C, followed by HRP-conjugated secondary antibodies for 2 h at room temperature. Signals were visualized using enhanced chemiluminescence and imaged with a cooled charge-coupled device system.

The primary antibodies used for immunoblotting were as follows: ADAMTSL4 Rabbit pAb (Abclonal, #A4785, 1:1000), TGFB2 Rabbit pAb (Abclonal, #A3640, 1:1000 dilution), Collagen VIII alpha 1 Rabbit pAb (Affinity, #DF8902,1:1000 dilution), GFP tag Mouse mAb (Proteintech, #66002-1-lg, 1:20000 dilution), mCherry Rabbit pAb (Proteintech, #26765-1-AP, 1:2000), and GAPDH Rabbit pAb (Proteintech, #10494-1-AP, 1:5000 dilution). The secondary antibodies used for immunoblotting were as follows: HRP-labeled Goat Anti-Mouse IgG(H+L) (Beyotime, #A0216, 1:1000 dilution) and HRP-labeled Goat Anti-Rabbit IgG(H+L) (Beyotime, #A0208, 1:1000 dilution).

### Statistical Analysis

Statistical analyses were performed using SPSS version 25.0 (64-bit edition, IBM Corp., Armonk, NY, USA). The normality of the data was confirmed by the Shapiro-Wilk test and the quantitative variables were presented as mean ± standard error and median (quartiles) where appropriate. The descriptive variables were shown in the counts or proportions. The student’s *t-*test or Mann–Whitney U test was applied as appropriate for comparing the two independent groups. The one-way analysis of variance or Kruskal-Wallis test was used to compare the three or more independent groups. All the analyses were 2-tailed, and the statistical significance was set as a *p*-value of < 0.05. The results of the two-sided tests were considered significant at *p* < 0.05.

## ACKNOWLEDGMENTS

We thank Dr. Jiatong Li and Mr. Han Long for their generous support. We are grateful to Prof. Wei Dai and her laboratory members for their technical assistance and helpful discussion.

## DATA AVAILABILITY STATEMENT

The datasets generated and/or analyzed during the present study are not publicly available but are available from the corresponding author upon reasonable request.

## ETHICS APPROVAL

The study was approved by the Human Research Ethics Committee of Fudan University.

## CONSENT TO PUBLISH DECLARATION

Not applicable

## AUTHOR CONTRIBUTIONS

All authors contributed to the study’s conception and design. Zexu Chen proposed this study, designed the study, and wrote the manuscript. The zebrafish models were constructed by Xinyi Huang. The clinical data were collected by Yongxiang Jiang, Qiuyi Huo, Min Zhang, Tianhui Chen, Zexu Chen, and Jiawan Nan. The immunostaining, immunoblotting, immunoprecipitation, and qPCR were performed by Wannan Jia, Yalei Wang, Xinyao Chen, and Zexu Chen. The cell lines were maintained and assessed by Wannan Jia. The phenotype profiling of zebrafish was conducted by Xinyi Huang. The RNA-seq and scRNA-seq data were analyzed by Xin Shen. Keji Jiang critically reviewed the manuscript. Yongxiang Jiang, Yan Pi, and Zexu Chen supervised the project. All authors read and approved the final manuscript.

---

**Supplementary Figure 1.**
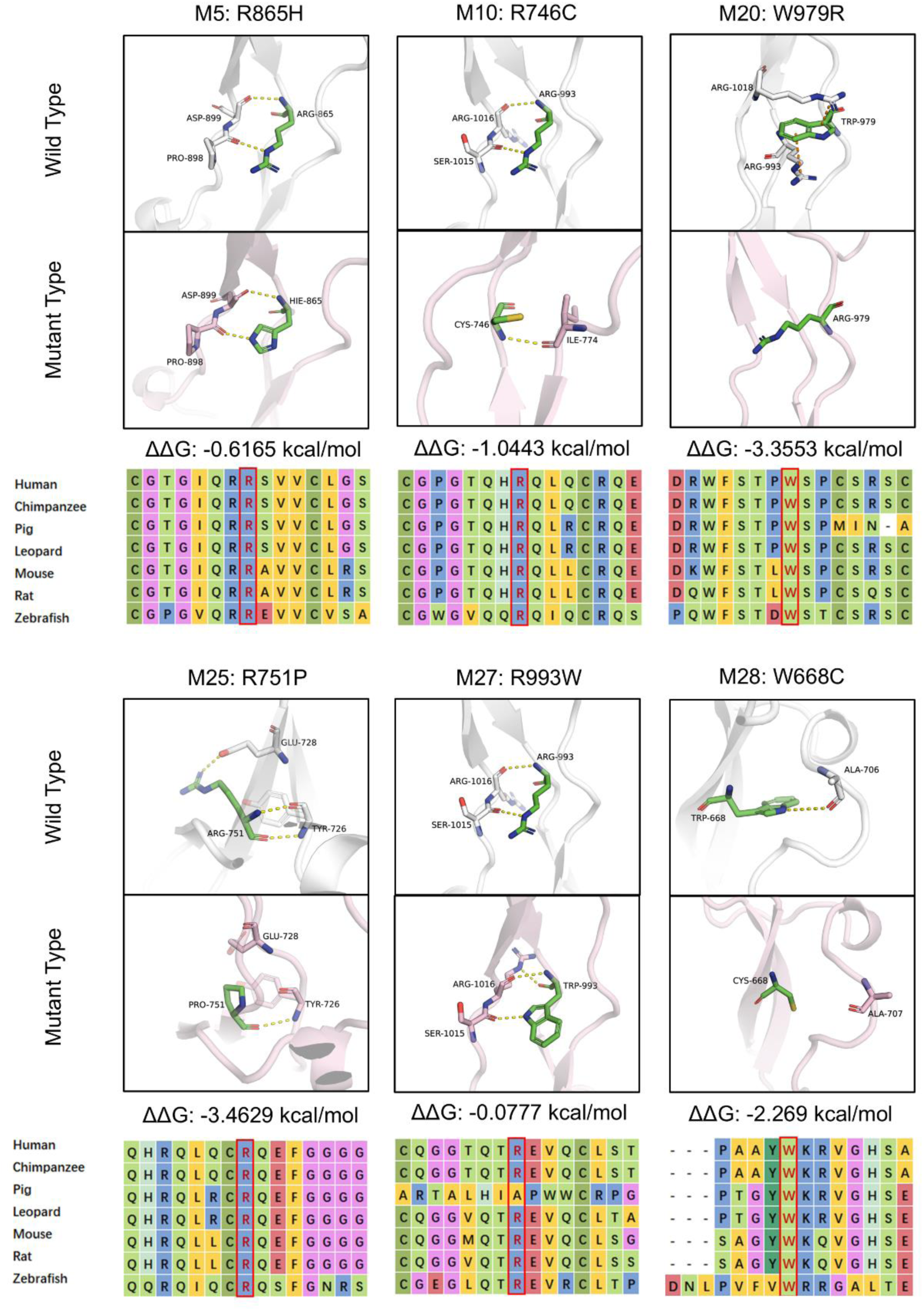
Structural Modeling and Multiple Sequence Alignment of *ADAMTSL4* Missense Variants Identified in This Study. The alteration of the protein stability was indicated by the free energy change value (ΔΔG = ΔG(mutant) -ΔG(wild-type), kcal/mol).

**Supplementary Figure 2.**
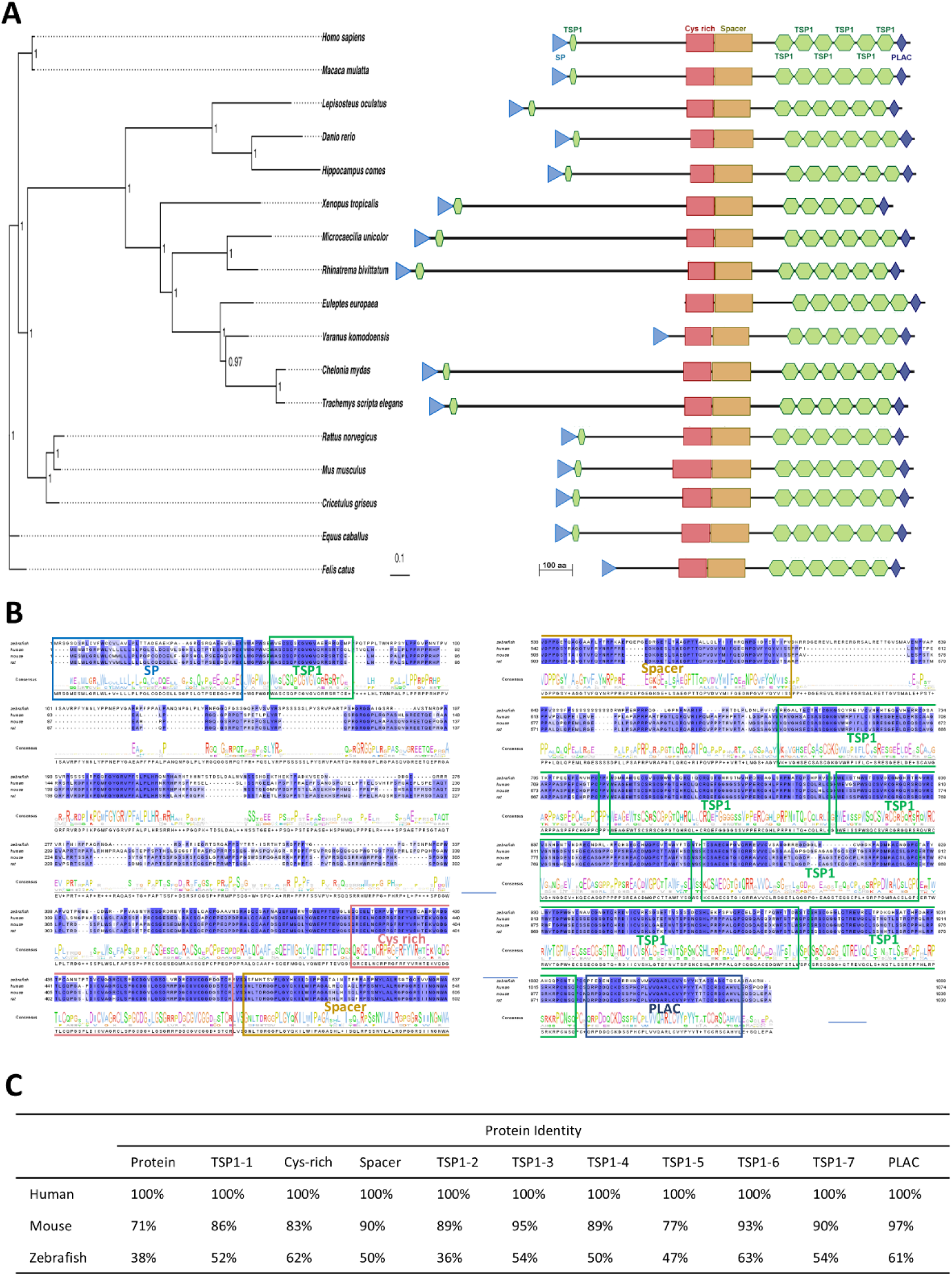
Evolutionary Conservation of *ADAMTSL4* in the Vertebrate Lineage. (A). Phylogenetic tree and protein domain diagrams illustrating the evolutionary relationship of *ADAMTSL4* orthologs in vertebrates. The tree was constructed using a best reciprocal BLAST search with human *ADAMTSL4*, mapping orthologs across diverse animal phyla. The tree is drawn to scale, with branch lengths representing evolutionary distances. Protein diagrams represent the longest isoform, highlighting conserved functional domains. Sequence data were obtained from the NCBI protein database. Scale bars: 0.10 (phylogenetic tree), 100 amino acids (protein diagram). (B). Protein sequence alignment of *ADAMTSL4* across zebrafish, human, mouse, and rat. Conserved domains are highlighted with boxes. (C). Protein identity and conserved domain comparisons among human, mouse, and zebrafish, calculated using reciprocal BLAST. aa, amino acid; BLAST, Basic Local Alignment Search Tool; NCBI, National Center for Biotechnology Information

**Supplementary Figure 3.**
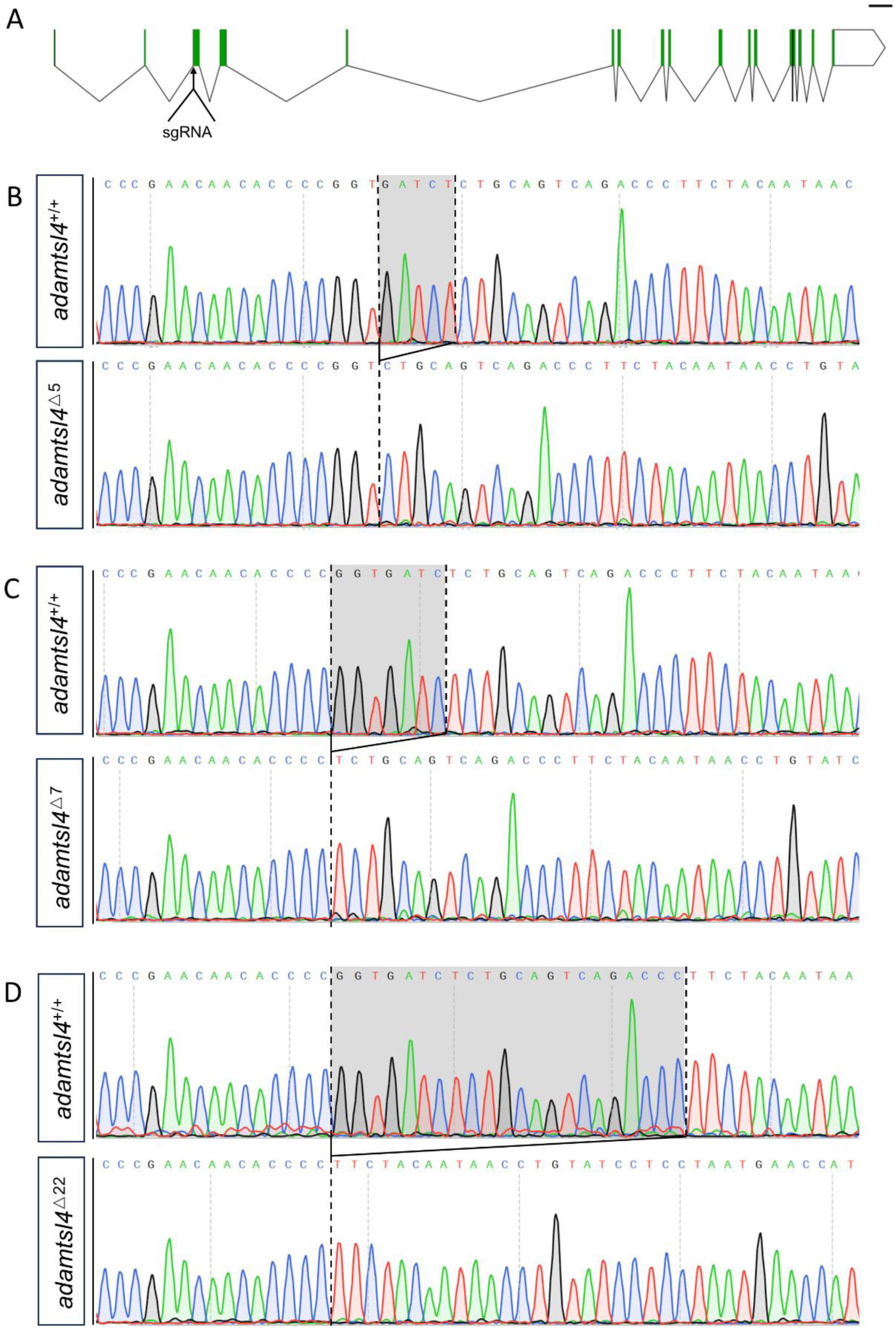
Sequence Validation of *adamtsl4* Knockout Mutants Generated by CRISPR/Cas9. (A). Schematic representation of the sgRNA target region used for CRISPR/Cas9-mediated knockout of *adamtsl4*. Scale bar, 2kb. (B). The *adamtsl4^△5^* mutant line validated by Sanger sequencing. (C). The *adamtsl4^△7^* mutant line validated by Sanger sequencing. (D). The *adamtsl4^△22^* mutant line validated by Sanger sequencing. sgRNA, single guide RNA

**Supplementary Figure 4.**
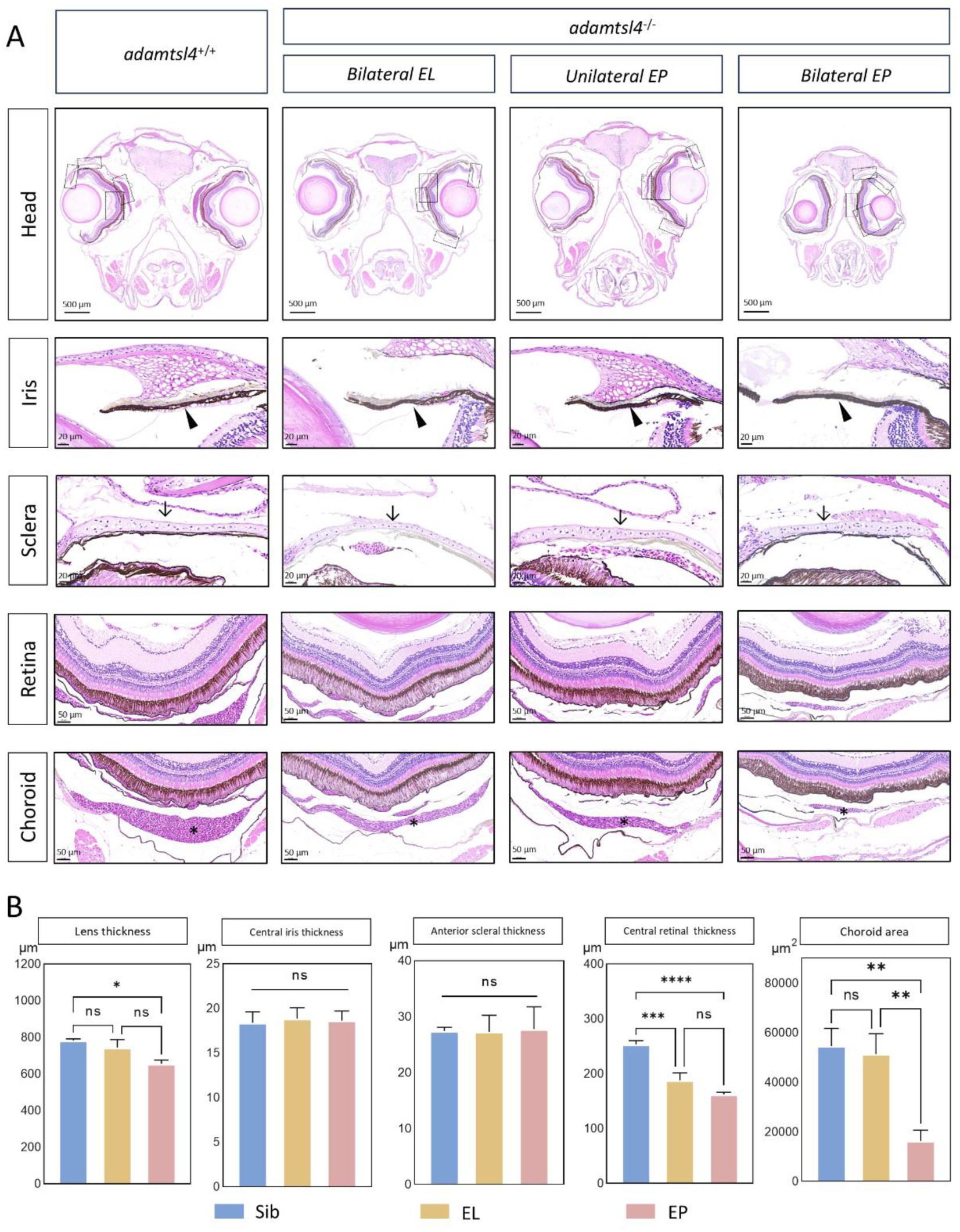
Histological Analysis of Ocular Structure of *adamtsl4^+/+^* and *adamtsl4^-/-^*Zebrafish. (A). Representative histological sections of adult zebrafish eyes, highlighting key ocular structures including the iris, sclera, retina, and choroid. Enlarged images of the boxed regions are provided for further detail. Scale bars: 500 μm (head), 50 μm (retina and choroid), 20 μm (iris and sclera). (B). Quantification of ocular parameters: lens thickness, central iris thickness, anterior sclera thickness, central retinal thickness, and choroid area. Sib refers to the normal eye of *adamtsl4^+/+^* sibling controls. EL refers to the "grossly normal" eye of *adamtsl4^−/−^* mutants, which exhibited ultrastructural zonular disorganization, whereas EP denotes the affected eye of *adamtsl4^−/−^* mutants with either unilateral or bilateral involvement. Statistical significance is indicated by asterisks: ns *P* ≥ 0.05, * *P* < 0.05, ** *P* < 0.01, *** *P* < 0.001, **** *P* < 0.0001; error bars indicate the standard error of the mean. EL, ectopia lentis; EP, ectopia pupillae; Sib, sibling control;

**Supplementary Figure 5.**
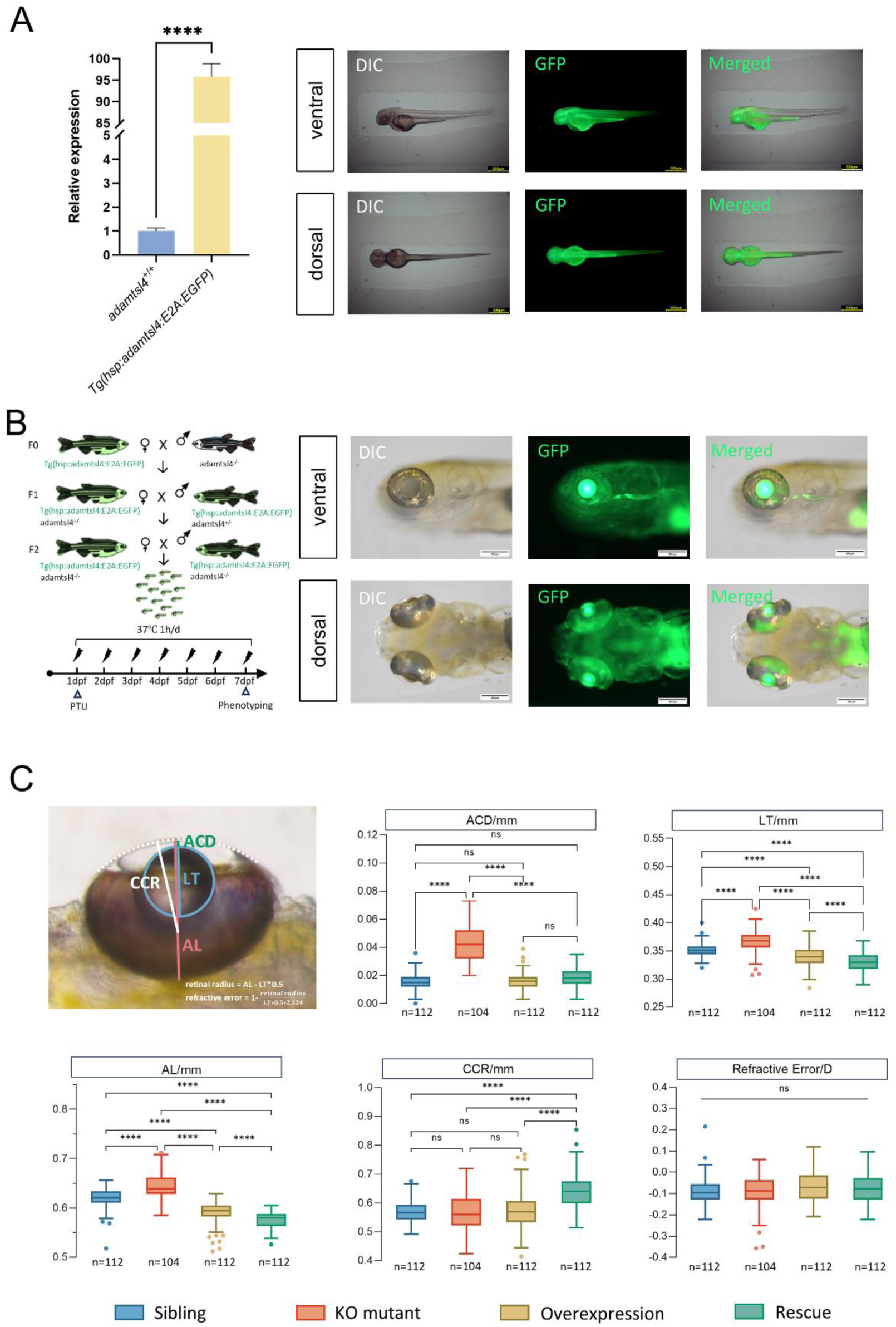
Overexpression and Rescue of *adamtsl4* in Zebrafish. (A). Validation of Tg(*hsp:adamtsl4:E2A:EGFP*) transgenic lines by qPCR and fluorescent imaging. Zebrafish larvae were heat-shocked at 37°C for 1 hour at 1 dpf. Scale bar = 500 mm. (B). Establishment of the rescue line. The Tg(*hsp:adamtsl4:E2A:EGFP*) transgenic was outcrossed with *adamtsl4^-/-^*lines and subsequently incrossed to generate a rescue line that carries both homozygous adamtsl4 mutations and the hsp-driven *adamtsl4* overexpression transgene. (C). Measurements of ACD, AL, CCR, LT, and refraction error in eyes of zebrafish at 7dpf. The groups compared include the sibling control (blue), *adamtsl4^-/-^*mutant (red), Tg(*hsp:adamtsl4:E2A:EGFP*) overexpression line (yellow), and rescue line (green). Statistical significance is indicated by asterisks: ns *P* ≥ 0.05, * *P* < 0.05, ** *P* < 0.01, *** *P* < 0.001, **** *P* < 0.0001; error bars indicate standard error of the mean; box plots display the median, quartiles, and variability of the data. ACD, anterior chamber depth; AL, axial length; CCR, corneal curvature radius; DIC, direct illumination contrast; dpf: day post-fertilization; KO, knock out; LT, lens thickness;

**Supplementary Figure 6.**
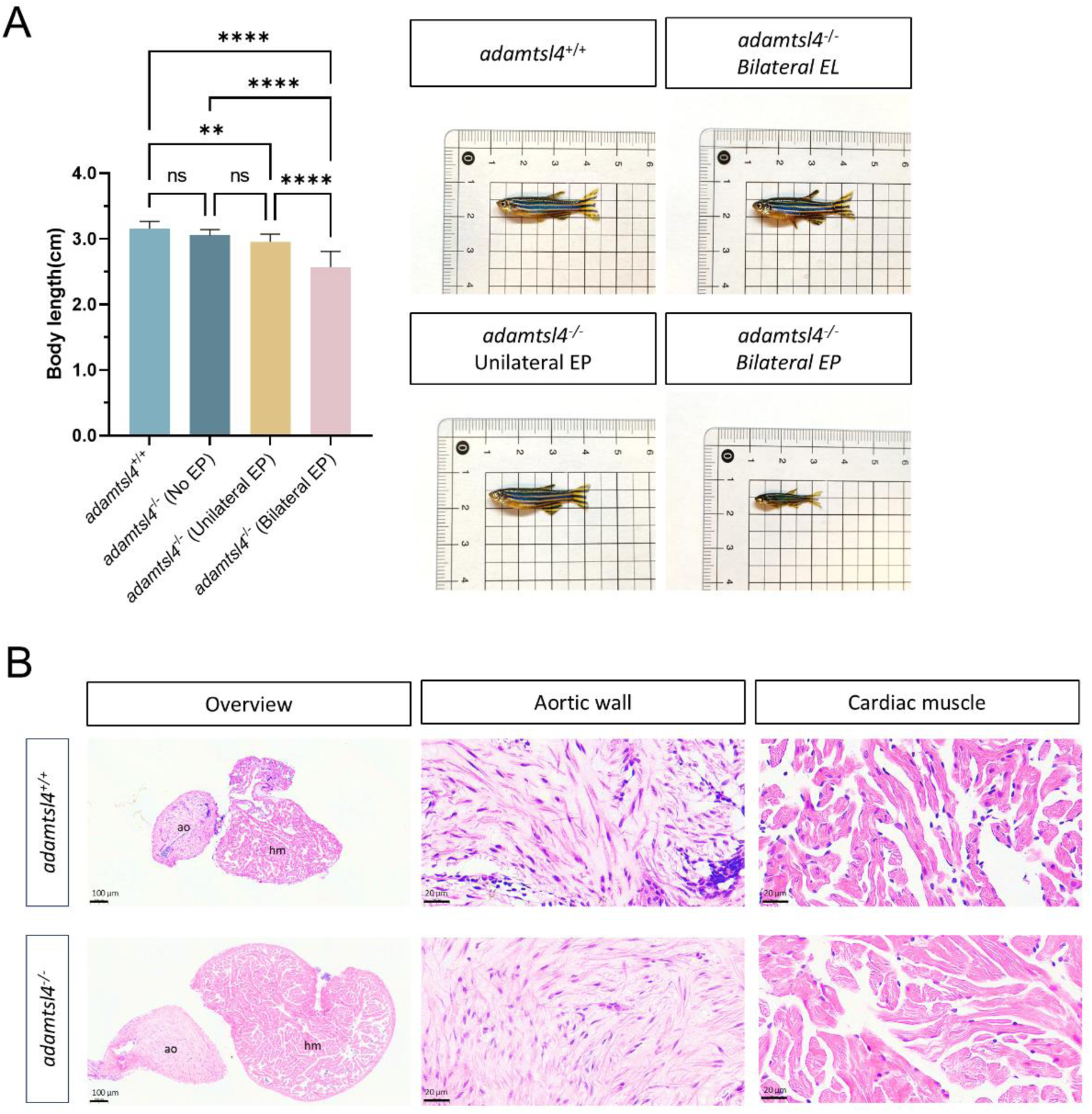
Systemic Phenotypic Features of *adamtsl4* Mutant Zebrafish. (A). Adult *adamtsl4^-/-^*zebrafish with EP exhibit retarded body size development compared to *adamtsl4^+/+^* sibling controls at 3 mpf. Each group consisted of 15 zebrafish. (B). Histological analysis of the heart and aorta phenotypes in *adamtsl4^-/-^* mutants and *adamtsl4^+/+^* sibling controls at 3 mpf. Statistical significance is indicated by asterisks: ns *P* ≥ 0.05, * *P* < 0.05, ** *P* < 0.01, *** P < 0.001, **** P < 0.0001; error bars indicate the standard error of the mean. ao, aorta; EL, ectopia lentis; EP, ectopia pupillae; hm, heart muscle; mpf, month post fertilization; UMAP, uniform manifold approximation and projection

**Supplementary Figure 7.**
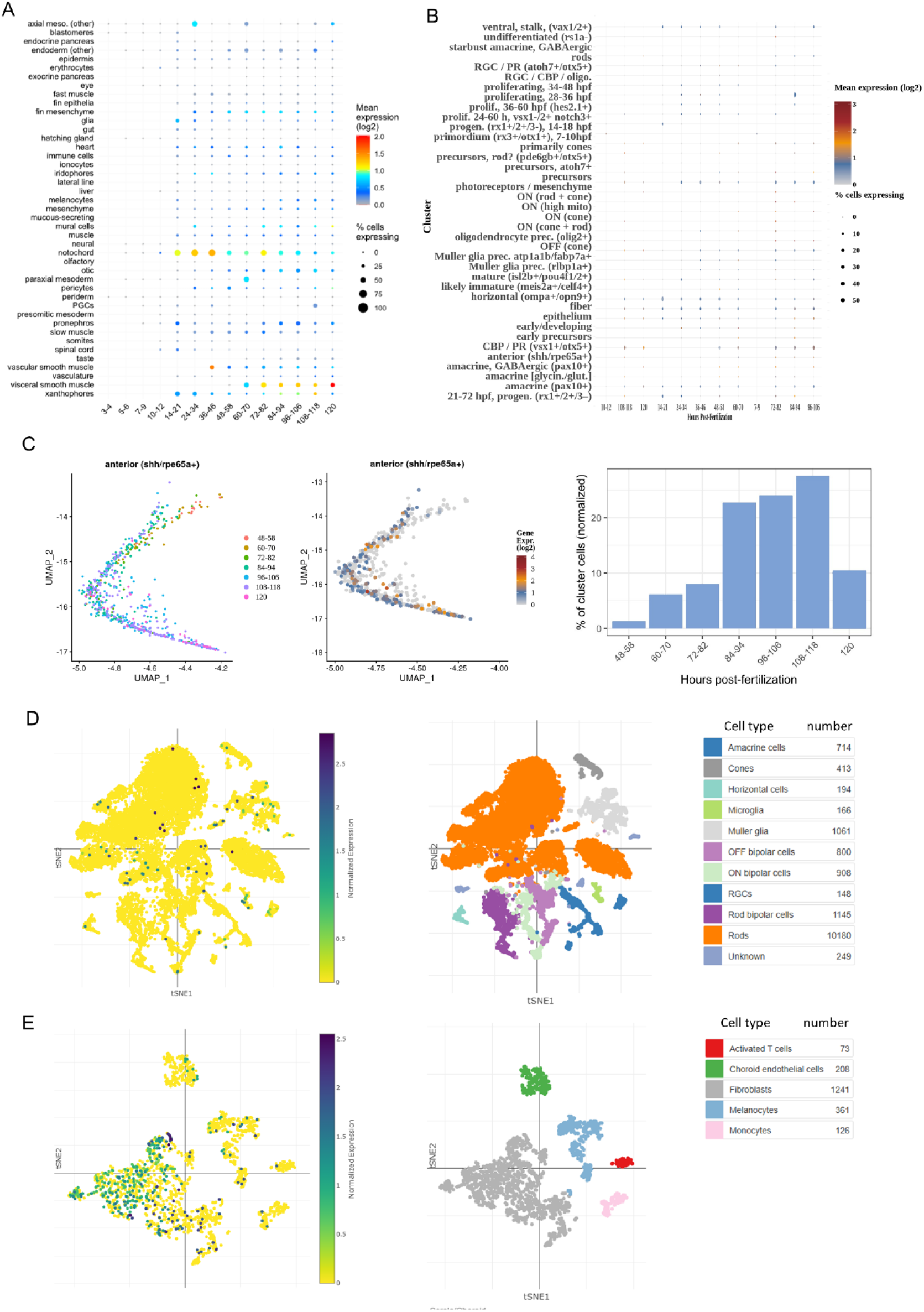
ScRNA-seq Reveals the Distribution Pattern of *ADAMTSL4/adamtsl4* Expression. (A). Temporal and spatial expression of *adamtsl4* in whole zebrafish larvae, showing expression across cell clusters and tissues. The size of each cluster represents the proportion of cells with *adamtsl4* expression > 0, while color intensity reflects the mean expression level within that cluster and timepoint. (B). Temporal and spatial expression of *adamtsl4* in ocular tissues of zebrafish larvae, with cluster size indicating the proportion of cells expressing *adamtsl4* > 0, and color indicating the mean expression level in cells from each cluster and timepoint. (C). UMAP embedding plot displaying *adamtsl4* expression in anterior RPE cells from 48 to 120 hpf. Color represents *adamtsl4* expression level, and clustering is based on developmental timepoint. The bar plot shows the percentage of anterior RPE cells from each developmental stage, normalized to the total number of cells collected at each stage. (D). tSNE embedding plot showing *ADAMTSL4* expression in the human retina. The expression level is indicated by color, with clustering based on different retinal cell types. (E). tSNE embedding plot showing *ADAMTSL4* expression in the human choroid. Color intensity represents expression levels, with clustering based on different choroidal cell types. PGC, primordial germ cell; RBC, retinal bipolar cell; RGC, retinal ganglion cell; RPE, retinal pigmented epithelium; prec., precursor; progen., progenitor; scRNA-seq, single-cell RNA sequencing; tSNE, t-distributed stochastic neighbor embedding; UMAP, uniform manifold approximation and projection.

